# Modular Arrangement of Synaptic and Intrinsic Homeostatic Plasticity within Visual Cortical Circuits

**DOI:** 10.1101/2024.06.01.596982

**Authors:** Wei Wen, Adriana M. Prada, Gina G. Turrigiano

## Abstract

Neocortical circuits use synaptic and intrinsic forms of homeostatic plasticity to stabilize key features of network activity, but whether these different homeostatic mechanisms act redundantly, or can be independently recruited to stabilize different network features, is unknown. Here we used pharmacological and genetic perturbations both *in vitro* and *in vivo* to determine whether synaptic scaling and intrinsic homeostatic plasticity (IHP) are arranged and recruited in a hierarchical or modular manner within L2/3 pyramidal neurons in rodent V1. Surprisingly, although the expression of synaptic scaling and IHP was dependent on overlapping signaling pathways, they could be independently recruited by manipulating spiking activity or NMDAR signaling, respectively. Further, we found that changes in visual experience that affect NMDAR activation but not mean firing selectively trigger IHP, without recruiting synaptic scaling. These findings support a modular model in which synaptic and intrinsic homeostatic plasticity respond to and stabilize distinct aspects of network activity.

## Introduction

To reliably maintain computational power in the face of experience-dependent modifications, neocortical networks use homeostatic forms of plasticity to constrain important features of network activity^1^. Defects in homeostatic plasticity contribute to pathological changes in network function by rendering circuits unable to compensate for perturbations arising during development or experience-dependent plasticity^2–8^. Many network features are under homeostatic control^9^, including mean firing rates^10,11^, sensory tuning curves^12,13^, nearness to criticality^14^, and the local correlation structure^15^. There is also a great diversity in the underlying cellular homeostatic plasticity mechanisms, but whether particular cellular mechanisms operate in a modular manner to regulate specific network features is still unknown^9^. For example, while computational work suggests that circuits may independently recruit synaptic and intrinsic homeostatic plasticity to regulate distinct aspects of network function^14–17^, recent experimental work has instead suggested that they are functionally and mechanistically coupled and are thus recruited in tandem^18^. Here we set out to determine whether synaptic and intrinsic homeostatic plasticity are arranged in a coupled, hierarchical manner, or whether they sense different features of network activity and thus can be independently recruited.

Circuit homeostasis is realized through careful adjustments of excitatory and inhibitory elements at a number of network nodes. Cellular mechanisms of homeostatic plasticity include those that modulate excitatory synapses, inhibitory synapses, or intrinsic neuronal excitability^9,19–21^. Among these, excitatory synaptic scaling and IHP have been observed in neocortical and hippocampal pyramidal neurons under a variety of experimental conditions^21–29^, and can be triggered together by global activity manipulations and sensory deprivation^27,29^. In the primary visual cortex (V1), monocular deprivation during the classic visual system critical period initially suppresses firing rates and reduces pairwise network correlations^8,11,15^; both of these network features are then homeostatically restored in parallel with the induction of synaptic scaling and IHP onto layer 2/3 (L2/3) pyramidal neurons^15,27^, and this homeostatic restoration is absent when these forms of plasticity are genetically ablated^8,29^. Despite their co-induction during sensory deprivation, it is still unclear whether synaptic scaling and IHP respond to the same features of altered activity, or whether they might be differentially sensitive to changes in firing rates and network correlations. A major factor in this information gap is the lack of knowledge about the induction and expression mechanisms of IHP, and the degree to which these are shared with synaptic scaling. For example, many of the classic activity manipulations used to study homeostatic plasticity, including blockade of spiking with tetrodotoxin (TTX), disrupt multiple calcium-dependent signaling pathways in parallel^9^, and which of these is necessary to trigger IHP is unknown.

To determine whether synaptic scaling and IHP share induction and expression mechanisms, and are arranged hierarchically or in a modular manner within neocortical circuits, we first propose two models (Figures 1A and 1B). In the first hierarchical scenario, the expression of synaptic scaling occurs prior to and results in the induction of IHP^18,30^ (Figure 1A), implying that they should always be triggered by the same activity manipulations. Alternatively, synaptic scaling and IHP could be arranged in a modular manner in which they sense changes in distinct aspects of activity (Figure 1B); in this model it is still possible that the two forms of plasticity converge onto shared trafficking pathways (upper panel).

**Figure 1.**
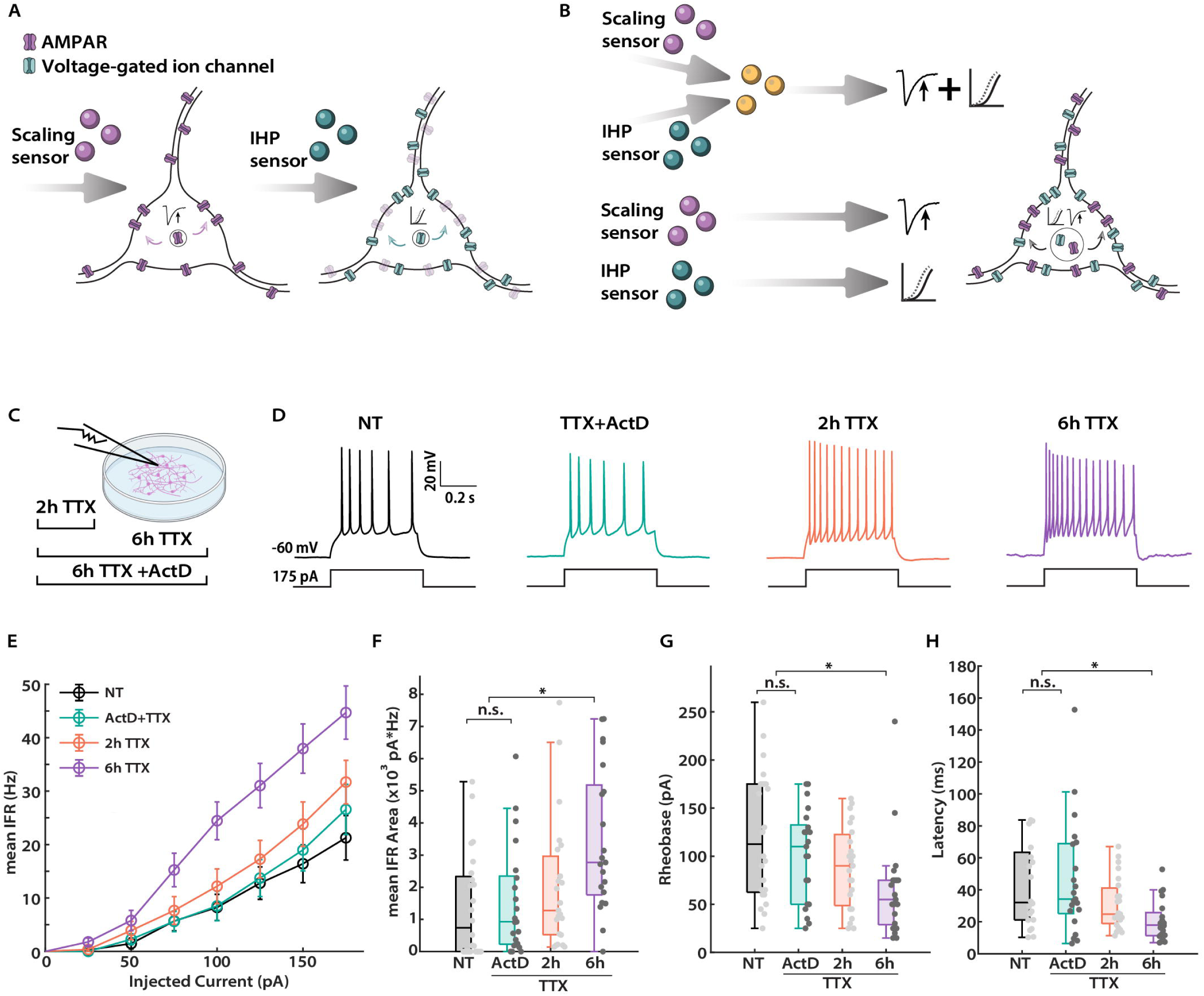
IHP is rapidly induced by TTX-mediated activity silencing and requires transcription. (A) and (B) Two proposed models for the coordination of synaptic scaling and IHP in neocortical circuits. (A) depicts a sequential model where synaptic scaling occurs prior to and is required for IHP induction. Alternatively, in (B), synaptic scaling and IHP are arranged in parallel and responsive to distinct activity sensors. Note that the intracellular processes controlling the expression of plasticity could converge (upper) or stay separated (lower). (C) Experimental paradigm. Cultured neocortical neurons were treated with TTX for either 2 or 6 hours before electrophysiological recording. To assess whether IHP expression requires gene transcription, neurons were also co-treated with TTX and Actinomycin D (ActD) for 6 hours. (D) Representative traces of spike trains evoked by current injections in cultured pyramidal neurons from the indicated conditions. (E) Comparison of mean instantaneous firing rate (IFR) vs. current injection (F-I) curves from the indicated conditions. NT, n = 24, 3 dissociations; ActD, n = 21, 3 dissociations; 2h, n = 23, 3 dissociations; 6h, n = 21, 3 dissociations. For F-I curves here and throughout, data are reported as mean±S.E.M of all cells from each condition. (F) Comparison of the area under the F-I curve for each neuron from the indicated conditions. Here and throughout, each data point indicates one neuron. Kruskal-Wallis test with Tukey correction: NT vs. ActD, p = 0.94; NT vs. 2h, p = 0.44; NT vs. 6h, p = 2.1E-3; ActD vs. 2h, p = 0.82; ActD vs. 6h, p = 0.02; 2h vs. 6h, p = 0.17. Here and below example comparisons are given on plots for clarity; * = p < 0.05, ** = p < 0.01, *** = p < 0.001, n.s. = not significant. (G) Comparison of rheobase current from the indicated conditions. Kruskal-Wallis test with Tukey correction: NT vs. ActD, p = 0.85; NT vs. 2h, p = 0.38; NT vs. 6h, p = 3.7E-3; ActD vs. 2h, p = 0.89; ActD vs. 6h, p = 0.06; 2h vs. 6h, p = 0.26. (H) Comparison of latency to the first spike for 175 pA of current injection from the indicated conditions. Kruskal-Wallis test with Tukey correction: NT vs. ActD, p = 1.0; NT vs. 2h, p = 0.71; NT vs. 6h, p = 0.01; ActD vs. 2h, p = 0.69; ActD vs. 6h, p = 0.01; 2h vs. 6h, p = 0.17.

To differentiate between these models, we perturbed different aspects of circuit activity *in vitro* and *in vivo*, and disrupted known trafficking pathways for synaptic scaling, to determine the co-dependence between synaptic scaling and IHP. We found that synaptic scaling and IHP share many aspects of their expression mechanisms, including time course, sensitivity to tumor necrosis factor α (TNFα), and intracellular trafficking pathways. Surprisingly, we also found that synaptic scaling and IHP are independently induced by reduced spiking and diminished NMDA receptor (NMDAR) signaling, respectively. Further, we found that a light-driven increase in pairwise correlations in V1 downregulates intrinsic excitability through an NMDAR-dependent mechanism, without inducing synaptic scaling. These data establish that synaptic scaling and IHP are driven by different activity sensors and are thus sensitive to distinct changes in circuit activity, yet rely on overlapping molecular pathways for their expression. Our results are consistent with a modular model in which synaptic scaling and IHP can be independently recruited to serve distinct functions within V1 circuits.

## Results

### IHP follows a similar expression timeline as synaptic scaling in vitro

If synaptic scaling and IHP share either activity-sensors or induction pathways, then they should follow a similar timecourse and be sensitive to the same molecular manipulations. In cultured cortical and hippocampal neurons, both synaptic scaling and IHP can be robustly induced by chronic activity silencing using TTX^23,28,31,32^. Further, synaptic scaling can be induced *in vitro* within 4 hours^33^, and by 6 hours is robust and almost as large in magnitude as at 24 hours^34^. To determine whether IHP follows a similar timecourse *in vitro*, we treated sister cultures with TTX for either 2 or 6 hours (Figure 1C), and measured the intrinsic excitability of pyramidal neurons by generating firing rate versus current (F-I) curves in the presence of synaptic blockers (Figures 1D and 1E). There was already an upward shift in the F-I curve after 2 hours of TTX treatment compared to the non-treated (NT) condition (Figure 1E, 2h TTX vs. NT), although the difference in the areas under the F-I curve was not yet statistically significant (Figure 1F, 2h TTX vs. NT). Intrinsic excitability increased significantly after 6 hours of TTX treatment, (Figures 1E and 1F, 6h TTX vs. NT), and was associated with lower rheobase current (Figure 1G), shorter latency to the first spike (Figure 1H), a subtle but significant narrowing of the first spike evoked at rheobase (Figure S1A, with no difference observed in the spike peak amplitude, Figure S1B), and higher neuronal input resistance (Figure S1C). In contrast, we did not observe any significant differences in action potential voltage threshold, spike frequency adaptation, resting membrane potential, or cell capacitance (Figures S1D-S1F, S5J). Therefore, IHP expression occurs rapidly in the first few hours of TTX treatment, closely following the timecourse of synaptic scaling.

A hallmark of TTX-induced synaptic scaling is its requirement for gene transcription^33,35–37^. To test whether IHP is also transcription-dependent, we co-treated cultures with actinomycin D (ActD) and TTX for 6 hours (Figures 1C and 1D). This prevented the normal TTX-induced change in the F-I curve (Figures 1E and 1F, ActD+TTX vs. NT), and there were no significant differences in other cellular and spiking properties (Figures 1G, 1H, S1A-S1F, S5J). Thus, like synaptic scaling, TTX-induced IHP depends on transcription in the early stage of its expression.

### IHP expression requires TNFα signaling

The expression of synaptic scaling is dependent on the cytokine TNFα, which maintains synapses in a plastic state that allows them to respond to perturbations in firing ^32,34,38^. Whether IHP is also dependent upon TNFα signaling is unknown. To test this, we began by assessing the dependence of IHP on TNFα in cultured cortical neurons, following the same experimental paradigm described in Steinmetz and Turrigiano^34^ (Figure S2A). An important finding of this study was that pretreating neurons with a TNFα scavenger^39^, sTNFR, for 24 hours completely blocked scaling up induced by 6 hours of TTX treatment. In contrast, co-treating neurons with sTNFR and TTX for 6 hours did not affect scaling up, indicating that a prolonged period of reduced TNFα signaling was necessary to block synaptic scaling. Intriguingly, the expression of TTX-induced IHP exhibited similar characteristics. While we still observed an increase in intrinsic excitability after 6 hours of TTX and sTNFR co-treatment (Figures S2B and S2C, sTNFR vs. Co), with the same suite of changes as described above for TTX alone (Figures S2D-S2F), pretreating these neurons with sTNFR for 14-17 hours before adding TTX prevented this increase in excitability (Figures S2B and S2C, sTNFR vs. Pre). These results demonstrate that – as for synaptic scaling – TNFα signaling plays a permissive role in TTX-induced IHP *in vitro*.

To determine whether this co-dependence on TNFα signaling is maintained in more mature *in vivo* circuits, we employed a previously validated activity suppression method, using the inhibitory DREADD hM4Di^29^ (Figure 2A). We targeted hM4Di to excitatory neurons in V1 by delivering a Cre-dependent hM4Di construct packaged in an adeno-associated viral vector into the V1 of Emx1-Cre mice during the classical visual system critical period. After waiting 7-10 days for robust DREADD expression, we administered CNO to the animals via drinking water for 24 hours to suppress V1 network activity^29^. In addition, we inhibited TNFα signaling *in vivo* by administering Xpro1595 (Xpro), a small-molecule TNFα scavenger that can pass the blood brain barrier and exerts measurable effects on plasticity for at least 48 hours after administration^38,40^. Based on our *in vitro* results, we pretreated animals with either Xpro or saline vehicle for 24 hours prior to the onset of the hM4Di silencing paradigm, followed by a second Xpro dose administered at the time animals were switched to CNO-containing drinking water (Figure 2A).

**Figure 2.**
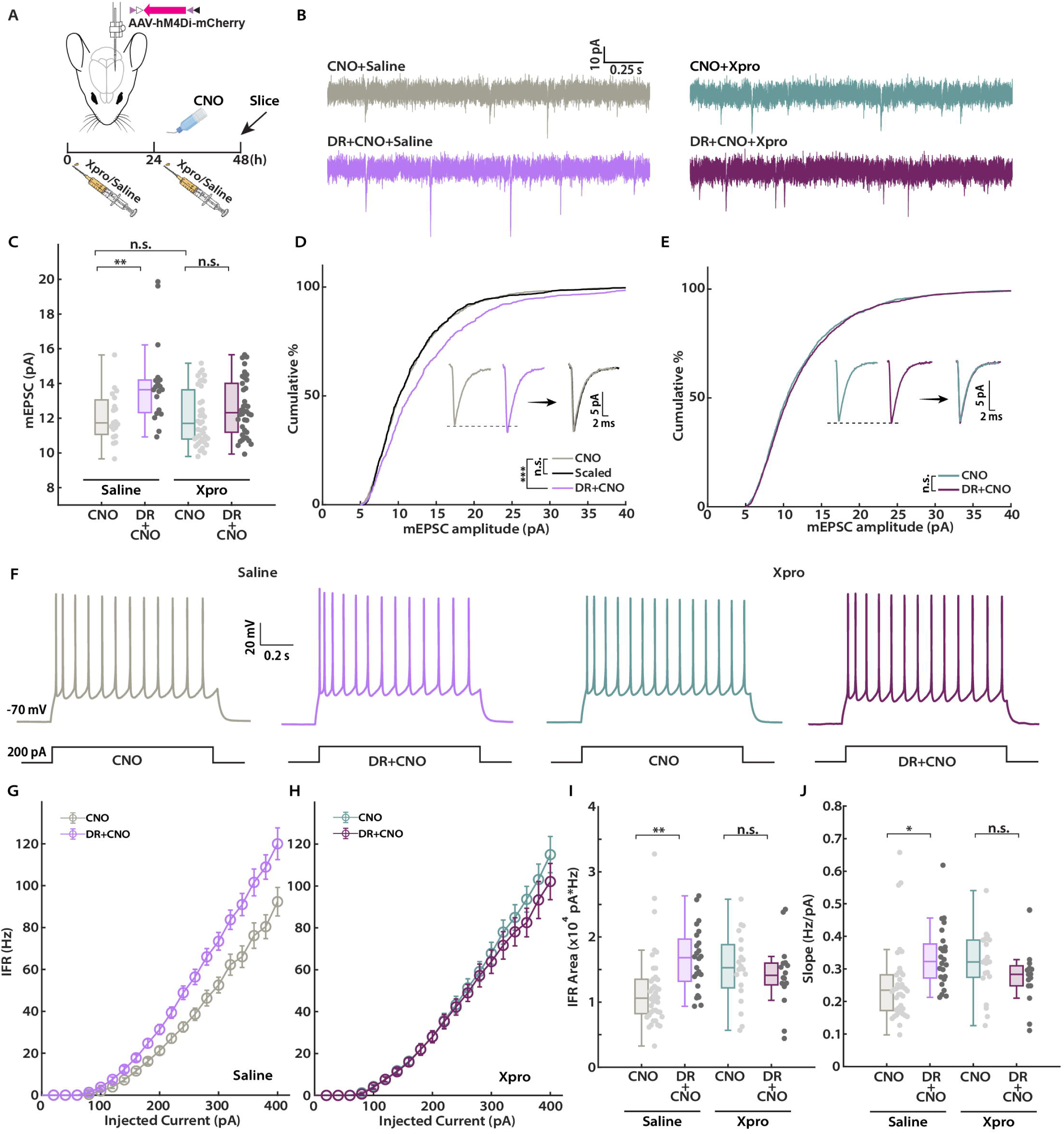
IHP expression in V1 L2/3 pyramidal neurons requires tumor necrosis factor α (TNFα) signaling. (A) Experimental paradigm. Viruses carrying hM4Di (DR) were unilaterally delivered into mouse V1 during P14-16. Animals were administered either Xpro or saline 24 hours before receiving CNO, and again when CNO was commenced. CNO was administered via drinking water for 24 hours before animals were sacrificed for slice electrophysiological recordings (during P25-30). Icons adapted from Biorender.com. (B) Representative traces of mEPSC recordings from the indicated conditions. (C) Comparison of the mean mEPSC amplitude for L2/3 pyramidal neurons from the indicated conditions. Saline, n = 19 from 6 animals for both CNO and DR+CNO conditions; Xpro, CNO, n = 39 from 9 animals, DR+CNO, n=38 from 9 animals. Two-way ANOVA test: interaction, p = 0.05; CNO vs. DR+CNO, p = 6.0E-4; Saline vs. Xpro: p = 0.04. Tukey correction: CNO+Saline vs. DR+CNO+Saline, p =5.6E-3; CNO+Saline vs. CNO+Xpro, p = 1.0; CNO+Saline vs. DR+CNO+Xpro, p = 0.73; DR+CNO+Saline vs. CNO+Xpro, p = 7.2E-4; DR+CNO+Saline vs. DR+CNO+Xpro, p = 0.03; CNO+Xpro vs. DR+CNO+Xpro, p = 0.53. (D) Cumulative distribution of mEPSC amplitudes for animals treated with saline. Population amplitudes from DR+CNO condition are scaled according to the linear function y = 1.35x – 2.21 (black curve). Inset: Left, unscaled average waveforms of CNO and DR+CNO conditions, respectively; Right, overlay of the average waveforms from CNO and scaled (black trace) populations, respectively. Kolmogorov-Smirnov test: CNO vs. DR+CNO, p = 9.1 E-12; CNO vs. Scaled, p = 0.56. (E) Same as D, but for animals treated with Xpro. Inset: Left, unscaled average waveforms of CNO and DR+CNO conditions, respectively; Right, overlay of the unscaled average waveforms from CNO and DR+CNO conditions, respectively. Kolmogorov-Smirnov test: CNO vs. DR+CNO, p = 0.09. (F) Representative traces of spike trains evoked by current injections in L2/3 pyramidal neurons from the indicated conditions. (G) Comparison of F-I curves for the CNO and DR+CNO groups from animals treated with saline. CNO, n = 40, 5 animals; DR+CNO, n = 24, 5 animals. (H) Same as G, but for animals treated with Xpro. CNO, n = 20, 4 animals; DR+CNO, n = 16, 4 animals. (I) Comparison of the area under F-I curve for each neuron from the indicated conditions. Two-way ANOVA test: interaction, p = 0.01; CNO vs. DR+CNO, p = 0.10; Saline vs. Xpro: p = 0.60. Tukey correction: CNO+Saline vs. DR+CNO+Saline, p =4.7E-3; CNO+Saline vs. CNO+Xpro, p = 0.09; CNO+Saline vs. DR+CNO+Xpro, p = 0.41; DR+CNO+Saline vs. CNO+Xpro, p = 0.87; DR+CNO+Saline vs. DR+CNO+Xpro, p = 0.55; CNO+Xpro vs. DR+CNO+Xpro, p = 0.94. (J) Comparison of the slope for the linear part of the F-I curve for each neuron from the indicated conditions. Two-way ANOVA test: interaction, p = 0.01; CNO vs. DR+CNO, p = 0.22; Saline vs. Xpro: p = 0.79. Tukey correction: CNO+Saline vs. DR+CNO+Saline, p =0.01; CNO+Saline vs. CNO+Xpro, p = 0.14; CNO+Saline vs. DR+CNO+Xpro, p = 0.71; DR+CNO+Saline vs. CNO+Xpro, p = 0.90; DR+CNO+Saline vs. DR+CNO+Xpro, p = 0.44; CNO+Xpro vs. DR+CNO+Xpro, p = 0.84.

First, we confirmed that Xpro blocks synaptic scaling. We cut slices of V1 and recorded miniature excitatory postsynaptic currents (mEPSCs) from L2/3 pyramidal neurons, and compared mEPSC amplitude after activity suppression combined with either Xpro or vehicle administration (Figure 2B). Comparing hM4Di-positive and hM4Di-negative neurons in the presence of saline or Xpro revealed a significant interaction (Two-way ANOVA, p = 0.05). There was a significant increase in the mean mEPSC amplitude in the presence of vehicle (Figure 2C, Saline, DR+CNO vs. CNO) and a clear rightward shift of the cumulative distribution of the entire mEPSC event population (Figure 2D), indicating synaptic scaling had occurred in these hM4Di-positive neurons, as expected^29^. In contrast, in the presence of Xpro mean mEPSC amplitude was unaffected by activity suppression (Figure 2C, Xpro, DR+CNO vs. CNO), and the cumulative amplitude distributions were similar (Figure 2E). Finally, Xpro had no significant impact on baseline mEPSC amplitudes (Figure 2C, CNO+Saline vs. CNO+Xpro). There were no significant differences in either the mean mEPSC frequency or waveform kinetics across conditions (Figures S2G-S2I, Insets of Figures 2D and 2E). Thus, inhibiting TNFα signaling is sufficient to block DREADD-induced synaptic scaling up in V1 L2/3 pyramidal neurons.

Next, we repeated the hM4Di-mediated silencing paradigm but instead measured F-I curves from L2/3 pyramidal neurons, in the presence of synaptic blockers (Figure 2F). As was the case for scaling, we observed a significant interaction between activity suppression and drug treatment (Two-way ANOVA, p = 0.01). Consistent with our previous results^29^, IHP expression was normal in saline-treated animals, indicated by comparable increases in both the area under F-I curve and the slope of the linear region (Figures 2G, 2I and 2J, Saline, DR+CNO vs. CNO). Further analyses of spike waveforms and trains revealed that this increase in intrinsic excitability in saline-treated animals was accompanied by reductions in the latency to the first spike (Figure S3H, Saline) and an increase in input resistance (Figure S3F, Saline). There was also a corresponding decrease in rheobase current, which did not reach statistical significance (Figure S3G, Saline). No significant differences were observed in the spike width at half-maximum (Figure S3B, Saline), the action potential voltage threshold (Figures S3A and S3C, Saline), the afterhyperpolarization amplitude (Figure S3D, Saline), spike peak amplitude (Figure S3E, Saline), spike frequency adaptation (Figure S3I, Saline), or in cell capacitance and resting membrane potential (Table S1). Strikingly, IHP was absent in Xpro-treated littermates, with no significant difference in the intrinsic excitability of hM4Di-negative and -positive neurons (Figures 2H,2I and 2J, Xpro, DR+CNO vs. CNO). In line with these findings, neurons from both conditions in Xpro-treated animals had no significant difference in any measures of excitability (Figures S3A-S3I, Xpro). Additionally, Xpro treatment did not significantly affect baseline intrinsic excitability (Figures 2I and 2J, CNO+Saline vs. CNO+Xpro). In conclusion, TNFα signaling is permissive for the expression of both synaptic scaling and IHP in the intact visual cortex.

### IHP is blocked by expression of a GluA2 c-terminal fragment in vivo

Our results so far establish that the induction of synaptic scaling and IHP share a number of features, suggesting that their induction might be interdependent. This could be explained by a hierarchical model, in which IHP is initiated by signaling pathways downstream of synaptic scaling expression (Figure 1A). This model predicts that blocking the expression of synaptic scaling will also abolish IHP.

To test this hypothesis, we manipulated a well-established molecular trafficking pathway for synaptic scaling to disrupt its expression. In neocortical and hippocampal pyramidal neurons, synaptic scaling relies on receptor trafficking interactions involving the GluA2 subunit of AMPA receptors (AMPARs)^35,41–43^, and can be blocked in L2/3 pyramidal neurons by the expression of a GluA2 C-terminal tail (C-tail) fragment^27,29^. To determine whether the GluA2 C-tail also blocks IHP, we virally co-expressed hM4Di and the GluA2 C-tail in one hemisphere of V1, while only expressing hM4Di in the other hemisphere as a positive control^29^ (CT+ and CT-, Figures 3A and 3B). After administering CNO for 24 hours, we measured intrinsic excitability of L2/3 hM4Di-positive pyramidal neurons (Figures 3C and 3D). While the F-I curve from CT-neurons shifted upward and leftward as expected (Figures 3D and 3E, CT- vs. CT+), there was no significant change in intrinsic excitability of CT+ neurons (Figure 3E, CT+ vs. NT). As expected, CT-neurons had a lower rheobase current (Figure 3F), a shorter latency to the first spike (Figure 3G), and a slightly higher neuronal input resistance (Figure 3H) than CT+ neurons. In summary, these results show that DREADD-induced IHP expression, like synaptic scaling, is susceptible to blockade by the GluA2 C-tail fragment.

**Figure 3.**
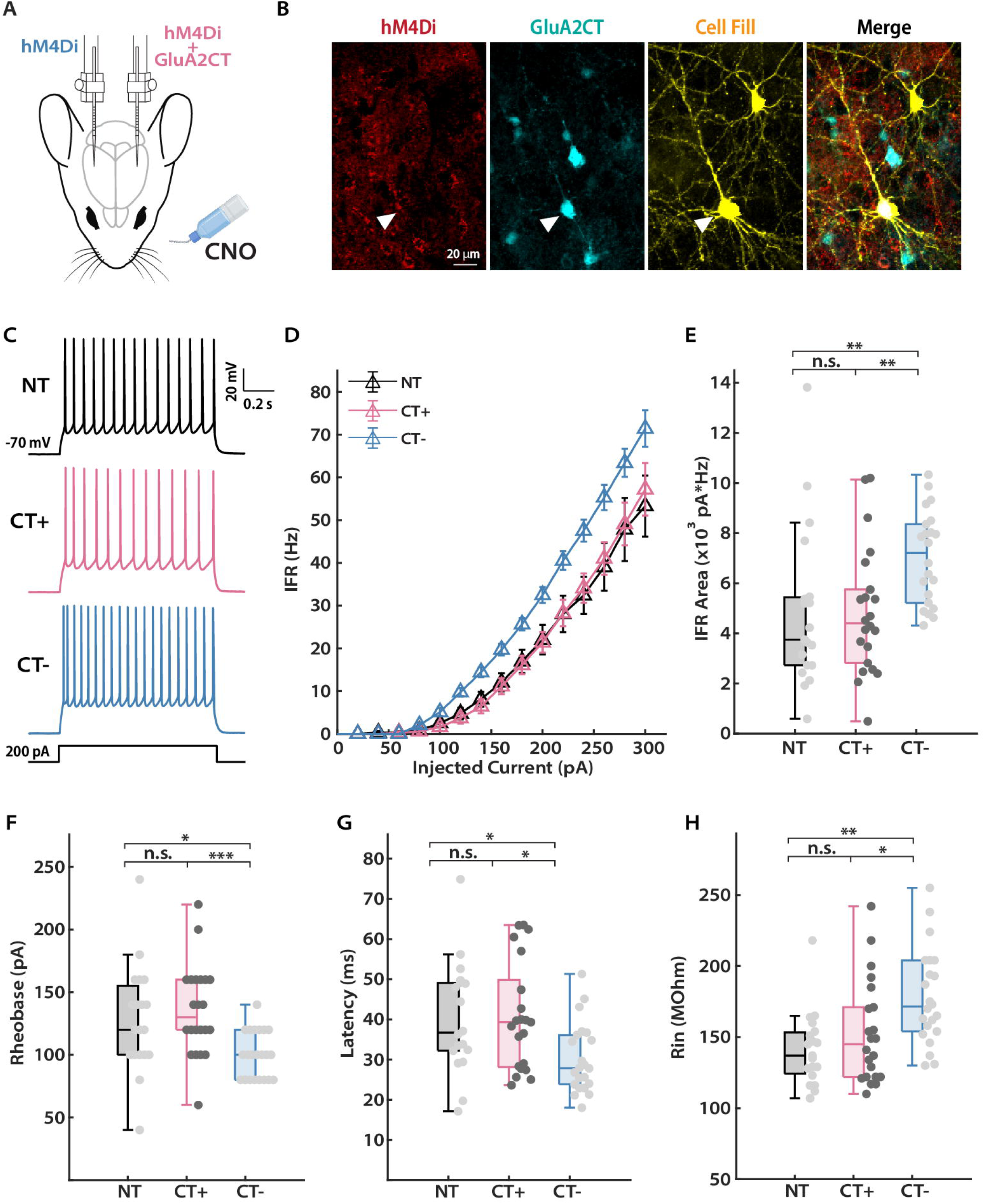
IHP in V1 L2/3 pyramidal neurons is prevented by expression of the GluA2 C-terminal fragment (CT). (A) Experimental paradigm. Viruses carrying the inhibitory DREADD hM4Di and GluA2 CT were mixed before being delivered unilaterally into mouse V1 during P14-16 (CT+), while only hM4Di was delivered into the other hemisphere to serve as a within-animal positive control (CT-). CNO was administered orally via drinking water for 24 hours before mice were sacrificed for slice electrophysiological recordings, which were performed during P25-30. Icons adapted from Biorender.com. (B) Representative images of a L2/3 pyramidal neurons recorded under the CT+ condition. The cell shows co-expression of hM4Di (tagged by mCherry) and GluA2 CT (tagged by GFP). White triangles indicate the recorded cell. Scale bar: 20 μm. (C) Representative traces of spike trains evoked by current injections in L2/3 pyramidal neurons for the indicated conditions. NT indicates littermates that received neither CNO nor virus injection. (D) Comparison of F-I curves of L2/3 pyramidal neurons from the indicated conditions. NT, n = 19, 3 animals; CT+, n = 22, 7 animals; CT-, n = 22, 7 animals. (E) Comparison of the area under F-I curve for each neuron from the indicated conditions. Kruskal-Wallis test with Tukey correction: NT vs. CT+, p = 0.95; NT vs. CT-, p = 4.7E-3; CT+ vs. CT-, p = 8.4E-3. (F) Comparison of rheobase current from the indicated conditions. Kruskal-Wallis test with Tukey correction: NT vs. CT+, p = 0.75; NT vs. CT-, p = 0.02; CT+ vs. CT-, p = 8.2E-4. (G) Comparison of latency to the first spike at 200 pA of current injection from the indicated conditions. Kruskal-Wallis test with Tukey correction: NT vs. CT+, p = 1.0; NT vs. CT-, p = 0.03; CT+ vs. CT-, p = 0.02. (H) Comparison of neuronal input resistance from the indicated conditions. Kruskal-Wallis test with Tukey correction: NT vs. CT+, p = 0.59; NT vs. CT-, p = 1.0E-3; CT+ vs. CT-, p = 0.02.

### Synaptic Scaling and IHP can be Independently Induced by suppressing spikes or NMDAR signaling, respectively

The sensitivity of IHP to manipulations that affect AMPAR trafficking is consistent with two different models for how synaptic scaling and IHP interact. First, as illustrated in the hierarchical model, IHP induction could be triggered by effectors downstream of synaptic scaling induction, such as calcium influx through glutamate receptors added during scaling^18^ (Figure 1A). In this scenario, IHP should not be inducible in the absence of synaptic scaling expression. Alternatively, synaptic scaling and IHP might be induced through separate signaling pathways, but rely on similar receptor/channel trafficking mechanisms during their expression (Figure 1B). To investigate these possibilities, we aimed to determine whether we could dissociate the induction of scaling and IHP by pharmacologically perturbing distinct aspects of circuit activity.

We started by treating cortical neurons from sister cultures either with TTX to block all spiking activity, or with an NMDAR antagonist, APV, to block NMDAR-mediated transmissions (Figure 4A). Importantly, TTX, which reduces but does not completely eliminate NMDAR activity^44,45^, robustly induces scaling^28^, whereas APV, which blocks NMDAR signaling but has little effect on spiking activity, does not induce scaling^28,46^. Previously we reported little effect of APV treatment on spontaneous firing rates in these neocortical cultures^28^; to confirm this we performed whole-cell recordings to measure spontaneous firing in pyramidal neurons in the presence and absence of acute APV, and found no significant difference in firing rates (Figures 4K and 4L). Further, we verified that mEPSC amplitudes were not increased after 24 hours of APV treatment, as expected^28^ (Figure 6E). To determine whether IHP is also insensitive to NMDAR blockade, we measured the intrinsic excitability of neurons treated for 24 hours with TTX, APV, or both (Figures 4B and 4C). Surprisingly, we found that APV treatment alone induced IHP with a comparable magnitude to that induced by TTX treatment (Figures 4C-4E, APV: 227% [area] and 148% [slope] of NT, TTX: 194% [area] and 167% [slope] of NT). Further, co-application of TTX and APV did not lead to a larger shift of the F-I curve (Figures 4C-4E, TTX+APV, 200% [area] and 154% [slope] of NT), indicating that these two manipulations are not additive. Last, the IHP expression observed under all three conditions was accompanied by similar changes in rheobase, spike latency, and input resistance (Figures 4F, S4F, S4G), suggesting that the expression mechanisms are similar. As in previous datasets we found no major difference in other cellular properties including action potential voltage threshold, spike width at half-maximum, spike peak amplitude, spike adaptation index, resting membrane potential, and cell capacitance (Figures S4A-S4E, S4H, S5J).

**Figure 4.**
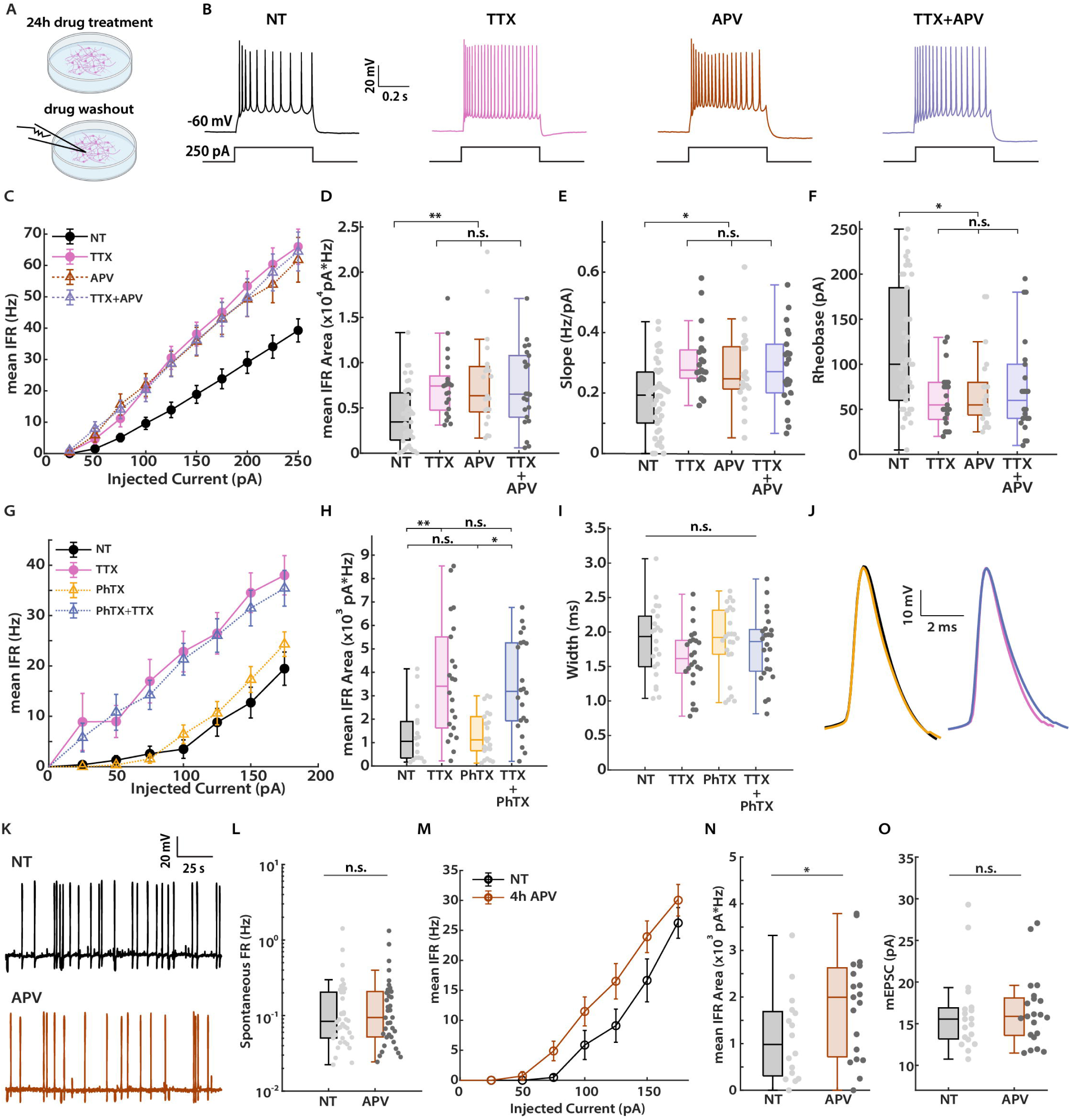
Synaptic scaling and IHP can be independently induced by suppressing spikes and NMDAR signaling, respectively. (A) Experimental paradigm for assessing effects of activity manipulation on homeostatic plasticity in cultured neocortical neurons. Electrophysiological recordings were performed after 24 hours of drug treatment. (B) Representative traces of spike trains evoked by current injections in cultured pyramidal neurons following treatments of TTX, APV, or TTX+APV. (C) Comparison of F-I curves for pyramidal neurons from the indicated conditions. NT, n = 42, 9 dissociations; TTX, n = 21, 5 dissociations; APV, n = 21, 6 dissociations; TTX+APV, n = 22, 3 dissociations. (D) Comparison of the area under F-I curve for each neuron from the indicated conditions. Kruskal-Wallis test with Tukey correction: NT vs. TTX, p = 3.7E-3; NT vs. APV, p = 0.01; NT vs. TTX+APV, p = 7.6E-3; TTX vs. APV, p = 0.99; TTX vs. TTX+APV, p = 1.0; APV vs. TTX+APV, p = 1.0. (E) Comparison of F-I curve slope from the indicated conditions. Kruskal-Wallis test with Tukey correction: NT vs. TTX, p = 1.0E-3; NT vs. APV, p = 0.03; NT vs. TTX+APV, p = 0.02; TTX vs. APV, p = 0.73; TTX vs. TTX+APV, p = 0.88; APV vs. TTX+APV, p = 0.99. (F) Comparison of rheobase current from the indicated conditions. Kruskal-Wallis test with Tukey correction: NT vs. TTX, p = 0.02; NT vs. APV, p = 0.04; NT vs. TTX+APV, p = 0.04; TTX vs. APV, p = 1.0; TTX vs. TTX+APV, p = 1.0; APV vs. TTX+APV, p = 1.0. (G) Comparison of F-I curves from the indicated conditions. NT, n = 22; PhTX, n = 24; TTX, n = 24; PhTX+TTX, n =24. (H) Comparison of the area under F-I curve for each neuron from the indicated conditions. Kruskal-Wallis test with Tukey correction: NT vs. PhTX, p = 0.83; NT vs. TTX, p = 1.1E-3; NT vs. PhTX+TTX, p = 1.4E-3; PhTX vs. TTX, p = 1.7E-3; PhTX vs. PhTX+TTX, p = 0.02; TTX vs. PhTX+TTX, p = 1.0. (I) Comparison of full width at half-maximum for the first spike at rheobase from the indicated conditions. Kruskal-Wallis test: p = 0.18. (J) Overlay of the average waveforms from first spikes evoked at rheobase. Left, NT vs. PhTX; Right, TTX vs. PhTX+TTX. (K) Representative traces of spontaneous firing recordings from cultured pyramidal neurons in the absence and presence of APV, respectively. NT, n = 38, 5 dissociations; APV, n = 37, 5 dissociations. (L) Comparison of spontaneous firing rate from the indicated conditions. Mann-Whitney U test: p = 0.71. (M) Comparison of F-I curves for pyramidal neurons from the indicated conditions. NT, n = 16, 3 dissociations; 4h APV, n = 19, 3 dissociations. (N) Comparison of the area under F-I curve for each neuron from the indicated conditions. Unpaired T test: p = 0.03. (O) Comparison of the mean mEPSC amplitude for pyramidal neurons from the indicated conditions. NT, n = 19, 3 dissociations; 4h APV, n = 21, 3 dissociations. Mann-Whitney U test: p = 0.64.

One study reported that short-term APV treatment can induce a transient increase in mEPSC amplitudes in cultured hippocampal neurons^44^. To rule out the possibility that synaptic scaling and IHP are transiently co-induced by short-term APV treatment, we measured both mEPSC amplitudes and F-I curves after 4 hours of APV treatment. While we observed a significant increase in intrinsic excitability (Figures 4M and 4N) that was similar in magnitude to that produced by short-term TTX treatment (Figures 1E and 1F), short-term APV treatment failed to increase mEPSC amplitude (Figure 4O). To rule out an even faster transient change we turned to direct measurement of synaptic AMPAR intensity after 1 hour of APV treatment, by quantifying the surface synaptic abundance of GluA2, and again found no significant change (Figures S4I and S4J). Thus IHP can be initiated independently of the expression of synaptic scaling.

It has been reported that a form of intrinsic homeostatic plasticity featuring action potential broadening can be triggered by calcium influx through calcium-permeable GluA1-containing AMPARs targeted to the synapses during chronic TTX silencing^18^. If activation of calcium-permeable AMPARs is critical for intrinsic plasticity, then blocking their activity should prevent IHP from occurring. We therefore treated cultured neurons with philanthotoxin (PhTX), a selective antagonist for calcium-permeable AMPARs, and assayed TTX-induced IHP (Figure 4G). Unlike TTX or APV, chronic PhTX treatment did not induce IHP on its own (Figures 4G and 4H, PhTX vs. NT). Furthermore, addition of PhTX did not prevent TTX-induced IHP (Figures 4G and 4H, PhTX+TTX vs. TTX), and there was no broadening of action potentials in any condition (Figures 4I and 4J), consistent with our previous datasets (Figures S1A, S4A and S4C). These results demonstrate that action potential broadening is not a feature of TTX- or APV-induced IHP in cortical pyramidal neurons measured either *in vitro* (Figure 4C) or *ex vivo* (Figure 2G), and activation of calcium-permeable AMPARs does not contribute to this form of IHP.

We next wished to test whether suppressing NMDAR activity alone *in vivo* would also induce IHP. We used CPP, an NMDAR antagonist that crosses the blood-brain barrier and produces a sustained reduction in NMDAR signaling after *in vivo* administration^47–50^, and blocks NMDAR-dependent Hebbian plasticity with little impact on visual function^49,51,52^. We used the same paradigm developed for the Xpro experiments, but instead of Xpro, we injected animals with CPP (Figure 5A). Following DREADD-mediated induction of IHP, we recorded F-I curves from L2/3 pyramidal neurons in the V1 of animals treated with either CPP or saline (Figures 5A-5C). Similar to the Xpro experiments, comparing all fours conditions revealed a significant interaction (Two-way ANOVA, p = 2.5E-3). Analyses of the F-I curves from these two groups of animals further revealed two clear effects (Figures 5C and 5D). First, neurons from CPP-treated animals exhibited higher intrinsic excitability than those from saline-treated littermates at baseline (Figure 5D, CNO+Saline vs. CNO+CPP). Second, despite the normal IHP expression in hM4Di-positive neurons from saline-treated animals (Figure 5D, Saline, DR+CNO vs. CNO), CPP induced no additional increase in intrinsic excitability in the hM4Di-positive neurons, indicating a lack of additivity (Figure 5D, CPP, DR+CNO vs. CNO, 95% of CNO). As for DREADD-mediated IHP, this CCP-induced increase in intrinsic excitability was associated with a lower rheobase, a shorter latency to the first spike, and a higher input resistance (Figures 5E-5G). These results show that CPP-mediated suppression of NMDAR activity induces IHP in the intact visual cortex, and likely occludes further DREADD-mediated IHP.

**Figure 5.**
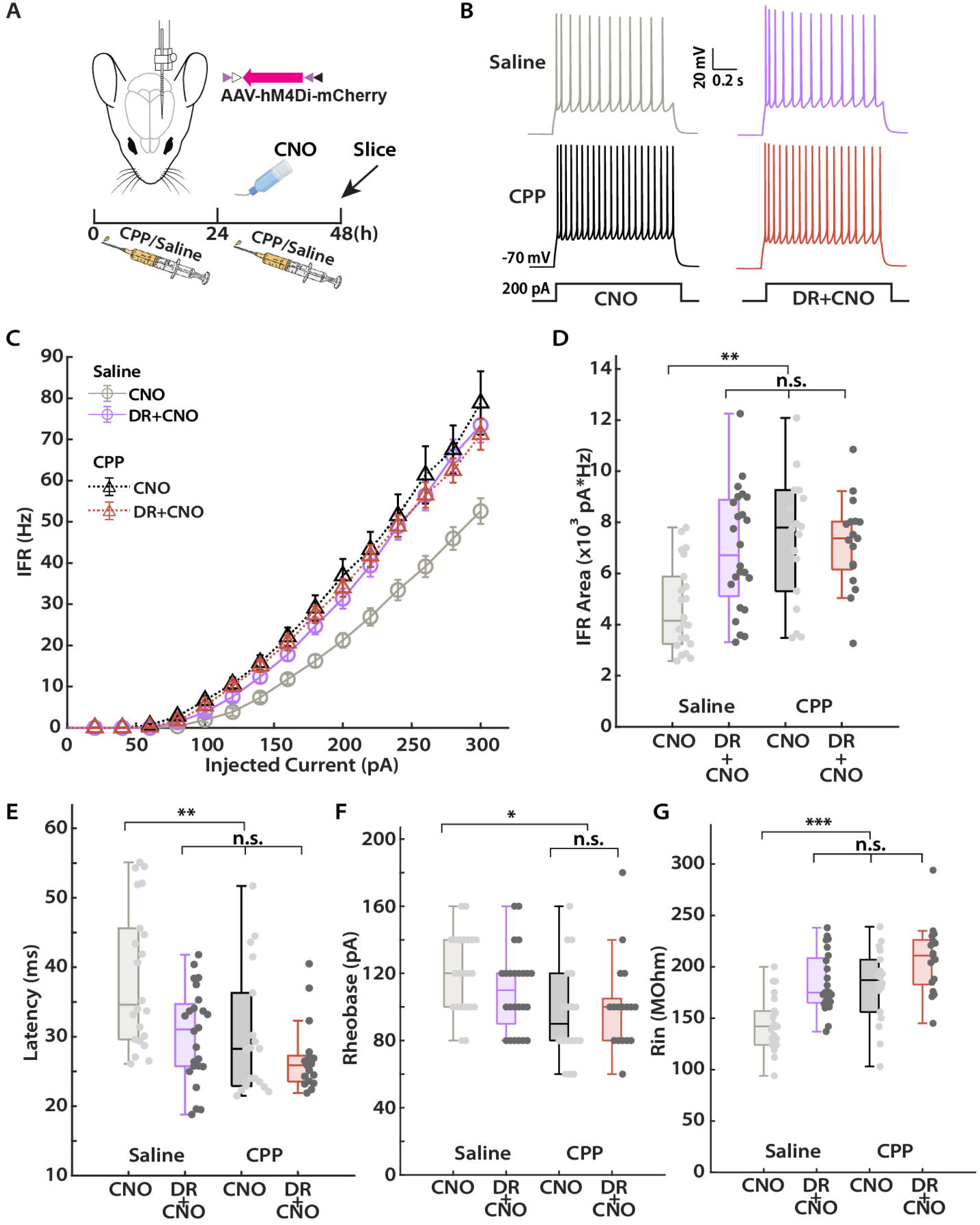
Chronic treatment with an NMDAR antagonist induces IHP in V1 L2/3 pyramidal neurons. (A) Experimental paradigm. Similar to the one performed in the Xpro treatment (Figure 2A), but with the NMDAR antagonist CPP instead. (B) Representative traces of spike trains evoked by current injections in L2/3 pyramidal neurons for the indicated conditions. (C) Comparison of F-I curves for L2/3 pyramidal neurons from the indicated conditions. Saline-CNO, n = 22, 5 animals; Saline-DR+CNO, n = 24, 5 animals; CPP-CNO, n = 18, 4 animals; CPP-DR+CNO, n = 17, 4 animals. (D) Comparison of the area under F-I curve for each neuron from the indicated conditions. Two-way ANOVA test: interaction, p = 2.5E-3; CNO vs. DR+CNO, p = 0.23; Saline vs. CPP: p = 9.2. E-3. Tukey correction: CNO+Saline vs. DR+CNO+Saline, p =7.7E-3; CNO+Saline vs. CNO+CPP, p = 6.0E-4; CNO+Saline vs. DR+CNO+CPP, p = 0.04; DR+CNO+Saline vs. CNO+CPP, p = 0.73; DR+CNO+Saline vs. DR+CNO+CPP, p = 0.99; CNO+CPP vs. DR+CNO+CPP, p = 0.59. (E) Comparison of latency to the first spike at 200 pA of current injection from the indicated conditions. Two-way ANOVA test: interaction, p = 0.12; CNO vs. DR+CNO, p = 1.5E-3; Saline vs. CPP: p = 1.6E-3. Tukey correction: CNO+Saline vs. DR+CNO+Saline, p =7.6E-4; CNO+Saline vs. CNO+CPP, p = 2.6E-3; CNO+Saline vs. DR+CNO+CPP, p = 3.8E-5; DR+CNO+Saline vs. CNO+CPP, p = 1.0; DR+CNO+Saline vs. DR+CNO+CPP, p = 0.68; CNO+CPP vs. DR+CNO+CPP, p = 0.72. (F) Comparison of rheobase current from the indicated conditions. Kruskal-Wallis test with Tukey correction: CNO+Saline vs. DR+CNO+Saline, p =0.48; CNO+Saline vs. CNO+CPP, p = 0.02; CNO+Saline vs. DR+CNO+CPP, p = 0.05; DR+CNO+Saline vs. CNO+CPP, p = 0.35; DR+CNO+Saline vs. DR+CNO+CPP, p = 0.61; CNO+CPP vs. DR+CNO+CPP, p = 0.98. (G) Comparison of neuronal input resistance from the indicated conditions. Two-way ANOVA test: interaction, p = 0.16; CNO vs. DR+CNO, p < 1E-324; Saline vs. CPP: p < 1E-324. Tukey correction: CNO+Saline vs. DR+CNO+Saline, p =4.8E-7; CNO+Saline vs. CNO+CPP, p = 1.5E-5; CNO+Saline vs. DR+CNO+CPP, p = 1.9E-11; DR+CNO+Saline vs. CNO+CPP, p = 0.99; DR+CNO+Saline vs. DR+CNO+CPP, p = 0.06; CNO+CPP vs. DR+CNO+CPP, p = 0.05.

Together these *in vitro* and *ex vivo* data provide strong evidence that IHP and synaptic scaling can be induced independently, and are sensitive to distinct aspects of neuronal activity: IHP is sensitive to reduced NMDAR signaling, while synaptic scaling is sensitive to reduced spiking and spike-mediated calcium influx^33^. These data argue against the hierarchical model (Figure 1A) and instead favor a model in which synaptic scaling and IHP are induced in parallel (Figure 1B).

### IHP, but not synaptic scaling, can be suppressed through allosteric modulation of NMDAR signaling

If IHP is triggered by reduced NMDAR signaling, then TTX should not be able to induce IHP under conditions where NMDAR activity is enhanced. To test this prediction, we used a positive allosteric modulator of NMDAR, GLYX-13 (GLYX)^53^, and examined its effect on TTX-induced IHP in cultured cortical neurons (Figure 6A). Strikingly, co-application of TTX and GLYX completely abolished IHP (Figures 6B and 6C, TTX+GLYX vs. NT), indicating that, when spiking is blocked, enhancing NMDAR activity is sufficient to prevent the induction of IHP. Examination of other cellular properties also revealed no difference between TTX +GLYX and the NT group (Figures S5A-S5F, S5J).

**Figure 6.**
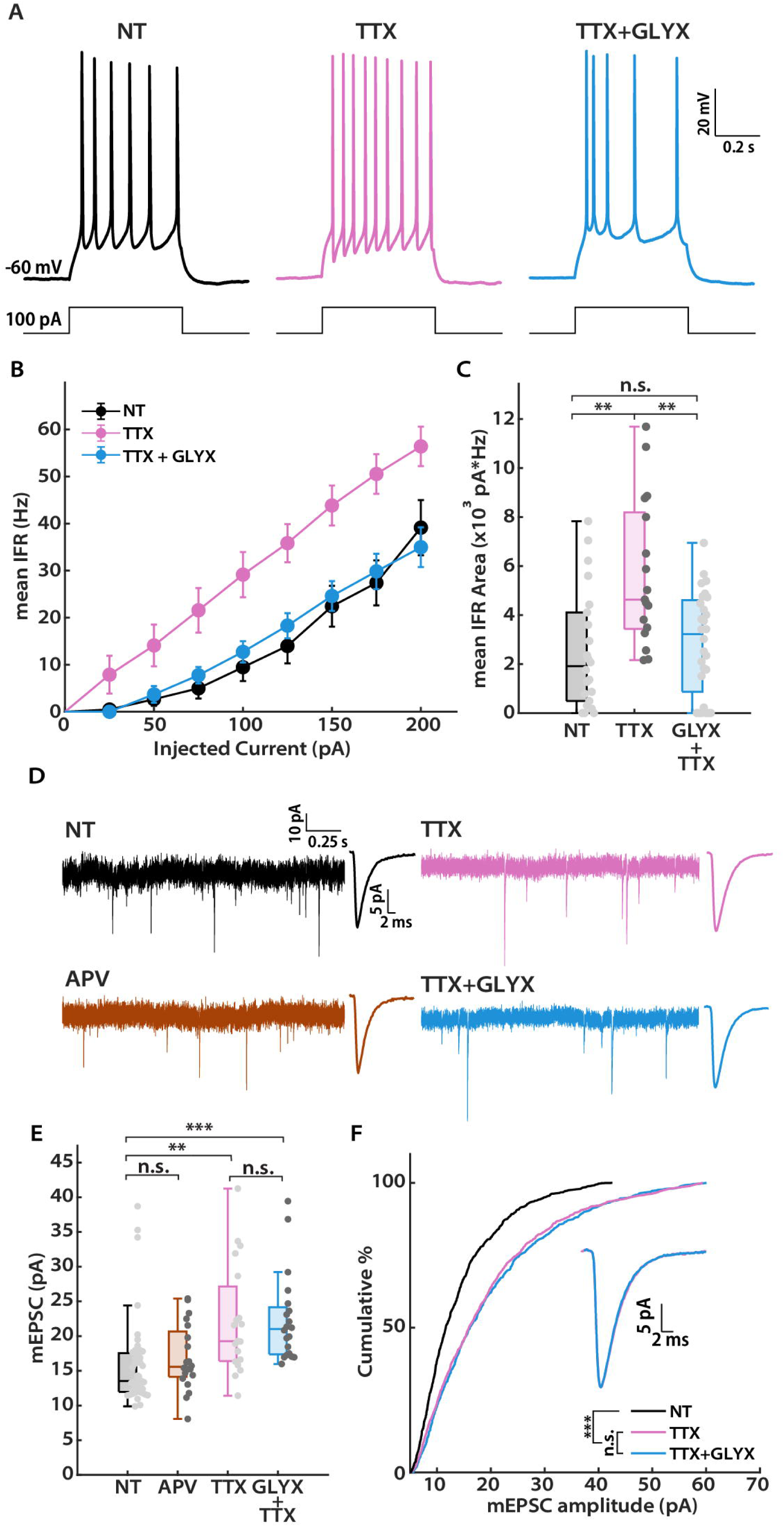
IHP, but not synaptic scaling, can be suppressed through enhancement of NMDAR signaling. (A) Representative traces of spike trains evoked by current injections in cultured pyramidal neurons from the indicated conditions. (B) Comparison of F-I curves for pyramidal neurons from the indicated conditions. NT, n = 22, 3 dissociations; TTX, n = 17, 3 dissociations; TTX+GLYX, n = 28, 3 dissociations. (C) Comparison of the area under F-I curve for each neuron from the indicated conditions. Kruskal-Wallis test with Tukey correction: NT vs. TTX, p = 0.01; NT vs. TTX+GLYX, p = 0.80; TTX vs. TTX+GLYX, p = 2.4E-3. (D) Representative traces of mEPSC recordings from the indicated conditions. The average waveform of mEPSC events is shown on the right of the corresponding recording trace. (E) Comparison of the mean mEPSC amplitude for pyramidal neurons from the indicated conditions. NT, n = 43, 6 dissociations; APV, n = 20, 3 dissociations; TTX, n = 19, 3 dissociations; TTX+GLYX, n = 20, 3 dissociations. Kruskal-Wallis test with Tukey correction: NT vs. APV, p = 0.62; NT vs. TTX, p = 1.5E-3; NT vs. TTX+GLYX, p = 3.4E-5; APV vs. TTX, p = 0.05; APV vs. TTX+GLYX, p = 0.02; TTX vs. TTX+GLYX, p = 0.90. (F) Cumulative distribution of mEPSC amplitudes. Inset: Overlay of unscaled average waveforms from TTX and TTX+GLYX conditions, respectively. Kolmogorov-Smirnov test: NT vs. TTX, p = 4.5E-42; NT vs. TTX+GLYX, p = 3.4E-64; TTX vs. TTX+GLYX, p = 0.17.

If NMDAR signaling is unimportant for the induction of synaptic scaling, then GLYX should have no impact on the ability of TTX to induce scaling up. To test this, we repeated the GLYX treatment and examined synaptic scaling by recording mEPSCs (Figure 6D). As expected, mEPSC amplitudes of neurons co-treated with GLYX and TTX were still scaled up when compared to the NT group (Figure 6E, TTX+GLYX vs. NT), to a similar degree as TTX alone (Figure 6E, TTX vs. NT). We then generated cumulative distributions of mEPSC amplitude from these conditions, and found that TTX and TTX+GLYX populations were statistically indistinguishable, but both were significantly different from the NT population (Figure 6F). GLYX had no effect on mEPSC frequency, waveform kinetics, or passive neuronal properties (Figures S5G-S5I). Thus, synaptic scaling will occur as long as spiking activity is suppressed. Finally, as described above, we verified that APV alone was not able to induce scaling up (Figure 6E, APV vs. NT). These data show that neither reducing nor enhancing NMDAR signaling influences synaptic scaling.

### Synaptic Scaling and IHP can be independently recruited during normal sensory experience

Taken together, the results described above show that IHP is induced via changes in NMDAR signaling, while synaptic scaling induction relies on altered spiking activity. An important implication of this finding is that it should be possible to independently recruit IHP without inducing synaptic scaling *in vivo* during experience-dependent manipulations that change NMDAR activation but leave mean firing rates unaffected. We showed previously that - while mean firing rates of V1 cortical neurons are stable across light and dark conditions - their pairwise correlations are significantly higher in the light than in the dark^54^. Given that correlated pre- and postsynaptic firing is a strong trigger for calcium influx through postsynaptic NMDARs^55^, we predicted that L2/3 pyramidal neurons would have higher intrinsic excitability in the dark (when correlations and thus NMDAR signaling is low) than in the light (when correlations and thus NMDAR signaling is high), while synaptic strength should be unaffected.

To test this idea, we first verified that pairwise correlations in V1 are higher in the light than in the dark, by re-analyzing chronic electrophysiological recordings of single-unit activity from V1 of freely behaving animals^51^. To follow the change in correlation across light and dark, we picked a 24-hour period spanning 12 hours of light and 12 hours of dark, and computed the pairwise correlation coefficients for all pairs of regular-spiking units (putative pyramidal neurons) in consecutive 30-minute bins. Plotting the mean normalized pairwise correlation against time clearly illustrated a higher average correlation in the light period, with a rapid drop in correlation at the transition to dark (Figure 7A). Comparing the correlation matrices from an ensemble of continuously recorded neurons within one animal across the light and dark periods revealed that the same pairwise correlations exist in both conditions, but are stronger in the light (Figure S6A). Comparing the normalized correlation for each pair of neurons demonstrates that the majority of these neuronal pairs exhibited higher correlation in the light (Figure S6B, 21/25 pairs). In contrast, across multiple V1 datasets collected under the same conditions mean firing rates remain stable across light and dark periods^10,11,54^.

**Figure 7.**
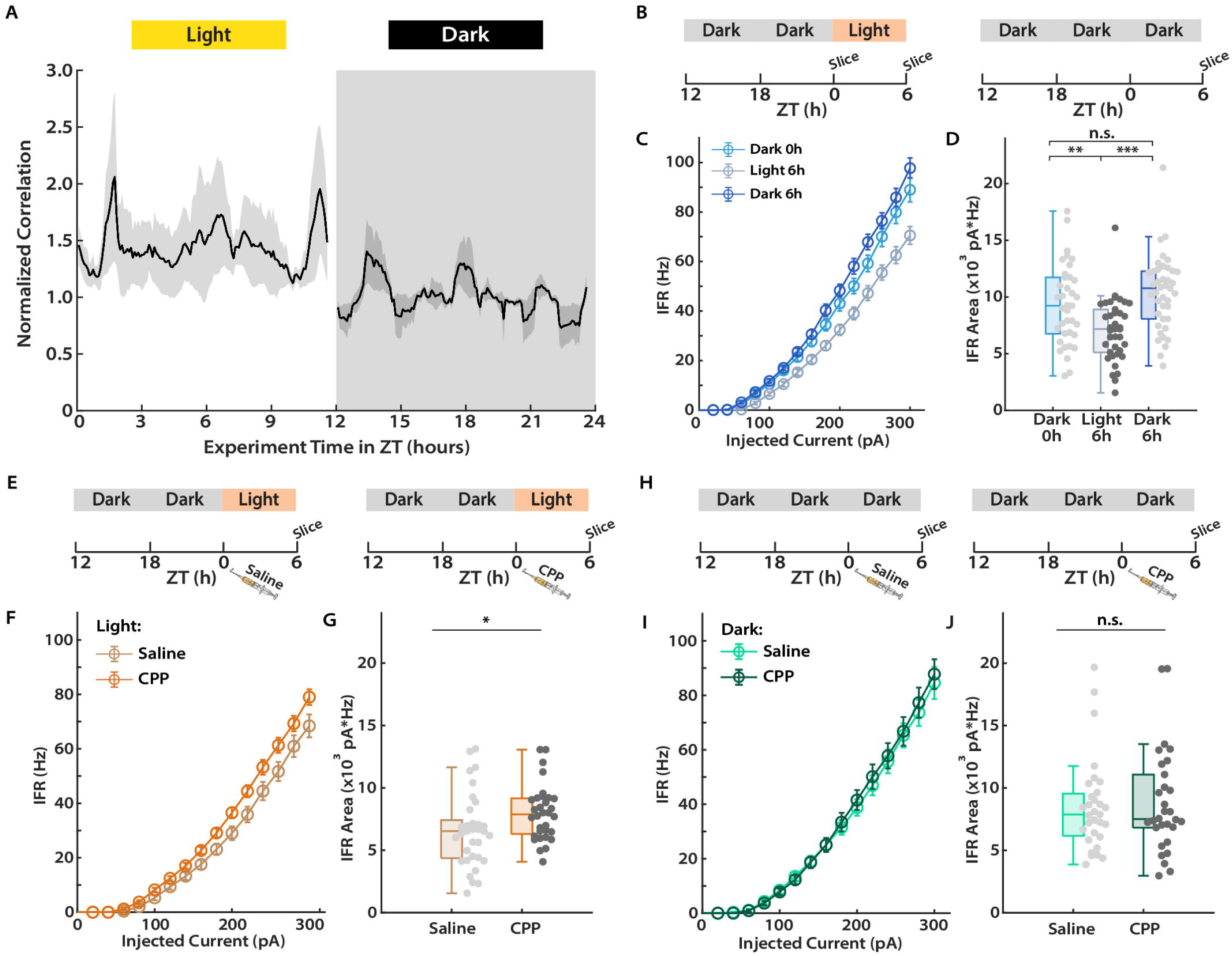
Synaptic scaling and IHP can be independently recruited during normal sensory experience. (A) The average correlation in firing for 25 pairs of regular spiking neurons from 5 animals over a day (12 hours light [white] + 12 hours dark [gray]). Correlation values for each pair are normalized to the mean correlation between zeitgeber time (ZT) 900min and ZT 1000 min (when fluctuations in correlation remain relatively low for over an hour). Shaded regions around the correlation trace indicate S.E.M. (B) Experimental paradigm for assessing intrinsic excitability of L2/3 pyramidal neurons in mouse V1 across light and dark. Littermates (during P27-32) were sacrificed at ZT 0h (Dark 0h) and ZT 6h (Light 6h) for electrophysiological recordings, respectively (Left). For assessing intrinsic excitability following prolonged dark exposure, animals were kept in dark for another 6 hours from ZT 0h (Dark 6h) before being sacrificed for electrophysiological recordings (Right). (C) Comparison of F-I curves for L2/3 pyramidal neurons from the three indicated timepoints. Dark 0h, n = 37, 4 animals; Light 6h, n = 38, 4 animals; Dark 6h, n = 41, 4 animals. (D) Comparison of the area under F-I curve for each neuron from the three indicated timepoints. One-way ANOVA test with Tukey post-hoc correction: Dark 0h vs. Light 6h, p = 3.6E-3; Dark 0h vs. Dark 6h, p = 0.76; Light 6h vs. Dark 6h, p = 2.3E-4. (E) Experimental paradigms for assessing effects of CPP on light-driven reduction in intrinsic excitability. Littermates received either CPP or saline injections at ZT 0h (just before light came on) and were sacrificed for electrophysiological recordings at ZT 6h (after spending 6 hours in the light). (F) Comparison of F-I curves for L2/3 pyramidal neurons from the two indicated conditions in I. CPP, n = 32, 3 animals; Saline, n = 33, 3 animals. (G) Comparison of the area under F-I curve for each neuron from the two indicated conditions. Unpaired T test, p = 0.02. (H) Comparison of F Experimental paradigms for assessing effects of CPP intrinsic excitability without light exposure. Littermates received either CPP or saline injections at ZT 0h (just before light came on) and were sacrificed for electrophysiological recordings at ZT 6h (after spending additional 6 hours in the dark). (I) F-I curves for L2/3 pyramidal neurons from the two indicated conditions in I, left panel. CPP, n = 30, 3 animals; Saline, n = 31, 3 animals. (J) Comparison of the area under F-I curve for each neuron from the two indicated conditions. Unpaired T test, p = 0.86.

If the light-driven increase in correlations enhances postsynaptic NMDAR activation, this should trigger a slow reduction in intrinsic excitability that should be robustly expressed after 6 hours (e.g. Figure 1F). To test this, we cut acute slices from littermates at two timepoints: right before the end of the 12-hour dark period (Figure 7B, left, zeitgeber time [ZT] 0h), and 6 hours into the subsequent light period (Figure 7B, left, ZT 6h). We then examined intrinsic excitability of V1 L2/3 pyramidal neurons at these two timepoints by recording F-I curves (Figure 7C). Indeed, intrinsic excitability was lower after animals spent 6 hours in the light than at the end of the dark period (Figures 7C and 7D, Light 6h vs. Dark 0h). To verify that this reduction in excitability was driven by light rather than circadian time, we kept a third group of littermates in the dark for an additional 6 hours before sacrificing them for recordings (Figure 7B, right, ZT 6h). Remarkably, when animals were kept in prolonged darkness there was no reduction in intrinsic excitability (Figures 7C and 7D, Dark 6h vs. Dark 0h), indicating that IHP is driven by light exposure rather than circadian entrainment. Further analyses of spike trains revealed that the light-driven reduction in intrinsic excitability was accompanied by increases in the rheobase (Figure S7A) and latency to first spike (Figure S7B), and a reduction in the neuronal input resistance (Figure S7C)-the same set of cellular changes that underlie DREADDs- and TTX-induced IHP induction. In sum, these results show that perturbations associated with light-driven changes in pairwise correlations trigger IHP in freely behaving animals.

Since the mean firing rates of V1 neurons are stable across light and dark, and we have shown that synaptic scaling is insensitive to changes in NMDAR signaling, mEPSC amplitude should be stable across the same light/dark conditions. To test this, we recorded mEPSCs from V1 L2/3 pyramidal neurons from acute slices obtained at the same timepoints (Figure S6E). As expected, there was no difference in either the mean mEPSC amplitude or the cumulative amplitude distributions across all three timepoints (Figures S6F and S6G). Furthermore, light/dark exposure had no impact on mEPSC kinetics (Figure S6G, inset; Figures S6H and S6I). On the other hand, we noticed a small increase in mEPSC frequency in the light condition that was absent in the prolonged dark condition (Figures S6J and S6K); this may reflect additional forms of plasticity set in motion by changes in correlation or other light-driven effects.

If light-driven correlations reduce intrinsic excitability by enhancing NMDAR activation, then this should be blocked or reduced by acutely suppressing NMDAR activity *in vivo* during light exposure. Further, the same manipulation should not affect intrinsic excitability if animals are kept in prolonged dark, when NMDAR signaling is already low. To test this, we first verified that acute CPP administration had no major effects on activity, by quantifying cFos expression in V1 2 hours after CPP or saline injection (Figure S6C); CPP had no significant impact on the percentage of cFos-positive neurons (Figure S6D, CPP vs. Saline). We then administered saline or CPP to littermates just before the dark to light transition, then sacrificed them for the assessment of intrinsic excitability 6 hours after lights on (Figure 7E, ZT 0h vs. ZT 6h). Consistent with our prediction, neurons from animals treated with CPP had higher intrinsic excitability than saline-treated littermates (Figures 7F and 7G), indicating that CPP-mediated NMDAR inhibition can partially reverse the light-driven reduction in intrinsic excitability. This CPP-induced increase in excitability was accompanied by the usual cellular changes associated with IHP induction (Figures S7E-S7G).

We then repeated this paradigm in a second cohort of animals, but kept them in prolonged dark for 6 hours after saline or CPP administration (Figure 7H, ZT 0h vs. ZT 6h). Again as predicted, there was no difference between the F-I curves recorded from the saline- and CPP-treated littermates after prolonged dark exposure (Figures 7I and 7J). Interestingly, although IHP induction is not associated with a change in capacitance (Figures S5J and S7D, Table S1), CPP treated animals had a small but significant reduction in cell capacitance regardless of whether the animals were exposed to light or dark (Figure S7H). This could reflect CPP-driven or circadian effects that do not contribute to IHP^56^. Together, these data demonstrate that the light-driven reduction in intrinsic excitability is realized through downstream signaling involving NMDARs. More broadly, our results make clear that synaptic scaling and IHP in L2/3 pyramidal neurons are driven by different activity-sensors, and are thus sensitive to distinct aspects of network activity.

## Discussion

Several forms of homeostatic plasticity have been documented within the same neocortical circuits, but the degree to which they act redundantly, or can be independently recruited to stabilize different aspects of network activity, is unclear. Here we used both *in vitro* and *ex vivo* electrophysiology paired with pharmacological and genetic manipulations to determine whether two such mechanisms, synaptic scaling and IHP, are arranged hierarchically or in parallel in neocortical pyramidal neurons. We found that synaptic scaling and IHP rely on many of the same molecular pathways for their expression, yet are sensitive to and activated by different features of network activity: while synaptic scaling is induced by changes in mean firing rates, IHP is instead induced by changes in NMDAR activation. Further, we demonstrated that sensory experience that alters NMDAR signaling but leaves mean firing rate unaffected selectively recruits IHP but not synaptic scaling. Our results show that synaptic scaling and IHP are triggered by different activity sensors, supporting a model where individual homeostatic plasticity mechanisms act in a modular manner to regulate distinct network features.

Despite a growing computational literature arguing that different cellular homeostatic mechanisms could be used to independently control important circuit features^14–17^, we have only a rudimentary understanding of when and how different forms of homeostatic plasticity are engaged within biological circuits. Like other forms of plasticity, homeostatic plasticity can be roughly split into induction and expression pathways (Figures 1A and 1B). During induction neurons detect a perturbation away from some activity set-point to initiate plasticity, while expression requires changes in ion channel surface abundance and distribution to generate a compensatory response. There is convincing evidence that synaptic scaling is induced when mean firing rate deviates from a firing rate set point^11,28,33,42^, and since synaptic scaling and IHP are often induced in parallel ^23,27–29^, it was reasonable to assume they are initiated by the same activity sensor during induction. One study compatible with this view proposed that a hierarchy exists between the induction of synaptic and intrinsic plasticity ^18^. In this scenario (Figure 1A), synaptic scaling occurs prior to IHP, enhances calcium influx through calcium-permeable AMPARs, and this then initiates a form of intrinsic plasticity that involves spike broadening. In contrast, our results clearly demonstrate that IHP can be induced independently of scaling; blocking NMDAR-mediated signaling *in vitro* or *in vivo* induces IHP but not synaptic scaling, IHP induction can be suppressed during spiking blockade by enhancing NMDAR signaling even though synaptic scaling still occurs, and IHP is not affected by blockade of calcium-permeable AMPARs. While it is possible that spike broadening occurs under some conditions via different induction rules, we found no evidence for spike broadening in any of our *in vitro* or *in vivo* manipulations that dramatically modulate activity, here (Figures S3B, 4I, S1A, S4C, S5D) or in previous work from our lab and others^29,57,58^. Importantly, our data showing that scaling and IHP can each be induced independently of the other rule out a hierarchical model, and instead establish that scaling and IHP are sensitive to distinct aspects of network activity (Figure 1B).

These results are surprising in light of many observations that scaling and IHP can be disrupted by the same molecular manipulations. For example, both are driven by changes in intracellular calcium^18,33,42^, are modulated by CaMKIV signaling^59^ (but see Trojanowski and Turrigiano^58^), and are sensitive to loss of the multidomain scaffolding protein Shank3^8,29^. Our findings strongly suggest that the calcium sources that trigger these two forms of plasticity are different; while scaling is more sensitive to spike-driven calcium influx^33^, we show here that IHP is likely dependent on calcium influx through NMDAR, consistent with previous studies^31,60^. Interestingly, while we see no impact of the competitive NMDAR antagonist APV on AMPAR-mediated transmission at either short or long time-scales, consistent with previous findings^61,62^, there is a growing literature showing that open-channel NMDAR antagonists such as ketamine and MK801 can rapidly potentiate AMPAR-mediated synaptic currents^63,64^. This unique effect of open-channel blockers may be attributed to their mechanism of action; recent evidence suggests that ketamine enhances AMPAR-mediated transmission not by modulating ion flow through the channel pore, but rather via structural changes that alter protein interactions in the postsynaptic density^65^. Our finding that neither short (1 or 4 hours) nor long (24 or 48 hours) APV treatment can induce synaptic scaling (Figure 4O, 6E, S4J) is consistent with previous reports^28,33,66,67^; thus, blocking NMDAR-mediated calcium influx using competitive antagonists does not potentiate AMPAR-mediated transmission, yet does trigger IHP. Taken together, this suggests that scaling and IHP are triggered via different calcium sources, but converge onto overlapping molecular pathways downstream of calcium influx during plasticity expression. This dependence on distinct calcium sources is an efficient way to allow synaptic scaling and IHP to sense different aspects of network activity, and thus serve distinct compensatory roles within the circuit.

While we have carefully quantified the impact of APV on NMDAR-mediated synaptic currents *in vitro* ^45^, it is much more difficult to precisely quantify this *in vivo,* so it remains unclear how big a change in NMDAR signaling is required to trigger IHP. Similarly, it is unclear how large a change in spiking activity is required to trigger scaling. We have shown that inhibitory DREADDs activation robustly reduces cFos expression in V1 and induces synaptic scaling^29^, while CPP has little effect on cFos expression and does not (Figure S6D), but because cFos is an indirect measure of activity, we cannot rule out that CPP has a small impact on firing rates that is not sufficient to induce synaptic scaling. One way to more directly assess the impact of CPP on V1 activity in the future would be to monitor population activity while locally applying competitive NMDAR antagonists, as has been done recently with non-competitive NMDAR antagonists in the primary somatosensory cortex^68^.

A variety of molecular players have been implicated in the expression of scaling^69,70^, but very little is known about the mechanisms underlying IHP. We show here that IHP expression shares many features and dependencies with synaptic scaling. Both require protein synthesis, are sensitive to TNFα signaling, involve trafficking mechanisms that are disrupted by expression of a GluA2 C-terminal fragment, and unfold with a similar time course. The endpoint of both synaptic scaling and IHP is the regulation of the surface abundance and distribution of the ion channels that determine postsynaptic strength (AMPAR) or intrinsic excitability (voltage-gated ion channels)^19,23,31^. It thus makes sense that both forms of plasticity utilize transcription- and translation-dependent protein synthesis^36,71–74^, and might share common protein trafficking and recycling/degradation pathways^35,37,43,70,75^. Synaptic scaling relies on C-terminal interactions between the GluA2 subunit of the AMPAR and synaptic proteins such as GRIP1^35,43^, and disruption of GluA2 trafficking via expression of a GluA2 C-tail fragment blocks scaling both *in vitro* and *in vivo* ^27,29,41^. Given that IHP does not require the prior induction of synaptic scaling, we were surprised to find that this same manipulation can also prevent IHP. One possible explanation is that this C-terminal fragment interferes with other more general ion channel trafficking pathways, such as the protein interactions with cell-adhesion and cytoskeletal molecules important for the membrane localization of voltage-gated ion channels^75,76^, or the recycling endosomal pathways that shuttle membrane-bound proteins between different membrane compartments^37,77,78^.

If synaptic scaling and IHP are functionally distinct, which circuit features do they stabilize, respectively? Many features of neocortical network activity are known to be under homeostatic control, including mean firing rates^10,11^, sensory tuning curves^12,13^, the nearness of the network to criticality^14^, and the local correlation structures^15^. The induction of synaptic scaling and IHP is correlated with the homeostatic recovery of mean firing rates and pairwise correlation structure during prolonged sensory deprivation^10,11,27^. Here we find that the induction of scaling is driven by changes in firing, while IHP is driven by changes in NMDAR signaling, which is modulated by correlated firing reflected in pairwise network correlations. Importantly, we found that light exposure, which drives an increase in the magnitude of existing pairwise correlations but has little impact on mean firing rates ^54^ (Figure 7A), results in a sufficient change in NMDAR-mediated signaling to trigger IHP, without triggering scaling. These findings show that neocortical circuits can selectively recruit IHP in response to changes in NMDAR signaling incurred through long-lasting changes in correlated firing, and potentially any other cellular or network changes that enhance NMDAR activation. Interestingly, IHP is strongly developmentally regulated, and in L2/3 pyramidal neurons is absent in adult animals^29^; consistent with this, a recent study found that input resistance in V1 L2/3 pyramidal neurons is not different in light vs dark^79^. It is unknown whether IHP persists in other cell types into adulthood, or whether its role in V1 is confined to an adolescent critical period.

Despite sensing correlations, it is unlikely that IHP feeds back to directly exert homeostatic control over pairwise correlation structure, as IHP develops gradually while the change in correlations across light and dark is abrupt and quite stable after the shift has occurred. Alternatively, because IHP is likely to modulate dendritic excitability^80–82^, it may be that IHP induction directly regulates NMDAR activation by affecting the ease with which synaptic inputs can depolarize the dendrite to remove the voltage-dependent magnesium block. In this scenario IHP would sense and homeostatically regulate NMDAR activation itself. Alternatively, IHP might regulates other circuit features such as the variance in the timing of spikes (coefficient of variation [CV] of interspike intervals)^11,17^. Indeed, we have consistently observed that IHP changes the slope of F-I curves rather than the firing threshold (Figures 2G, S3C, 4E, S4B), which is predicted to magnify the impact of both excitatory and inhibitory inputs on spike probability, thus magnifying spike variance; IHP induction during monocular deprivation would then restore CV, as observed experimentally^11^. Either of these scenarios would leave synaptic scaling free to homeostatically regulate mean firing rate and/or pairwise correlation structure.

It has long been proposed that different forms of cellular homeostatic plasticity are instrumental in the homeostatic maintenance of distinct features of network activity, yet how homeostatic plasticity is arranged within neural circuits to achieve this was unclear. Taken together our results demonstrate that synaptic and intrinsic forms of homeostatic plasticity, even when expressed in the same cell type, sense distinct aspects of activity perturbations and thus can be independently recruited *in vivo*. These mechanistic differences between synaptic and intrinsic plasticity induction allows them to act either in concert or independently, and to regulate distinct aspects of network function. Together, our findings support the idea that key network features are independently constrained by different cellular homeostatic mechanisms to ensure that circuits can remain functional in the face of a wide range of perturbations.

### Limitations of the study

This study relies on pharmacological manipulations to selectively manipulate NMDAR and TNFα signaling; such manipulations may have incomplete specificity, and the *in vivo* pharmacokinetics of these drugs are incompletely understood. Further, we used a cFos-based detection method to assess the effect of CPP-mediated NMDAR blockade on circuit activity; while this is a well-established approach to detect bidirectional activity changes, it likely does not have the resolution to detect subtle differences in activity. A conceptual limitation is that we confine our analysis to one excitatory cell type within V1, so the generality of our conclusions about the induction requirements for synaptic scaling and IHP remain to be determined.

## Acknowledgments

We thank members of the Turrigiano lab for constructive discussions, Lirong Wang for help with neuronal culture maintenance, and the Brandeis Light Microscopy Core Facility (RRID:SCR_025892). This work was supported by R35NS111562 (G.G.T.). The graphic abstract was created with Biorender.

## Author Contributions

Conceptualization, W.W. and G.G.T.; Methodology, W.W., A.M.P., and G.G.T.; Investigation, W.W. and A.M.P.; Writing- Original Draft, W.W.; Writing- Review & Editing, W.W., A.M.P., and G.G.T.; Visualization, W.W and A.M.P; Funding Acquisition, G.G.T.

## Methods

### Key resource table

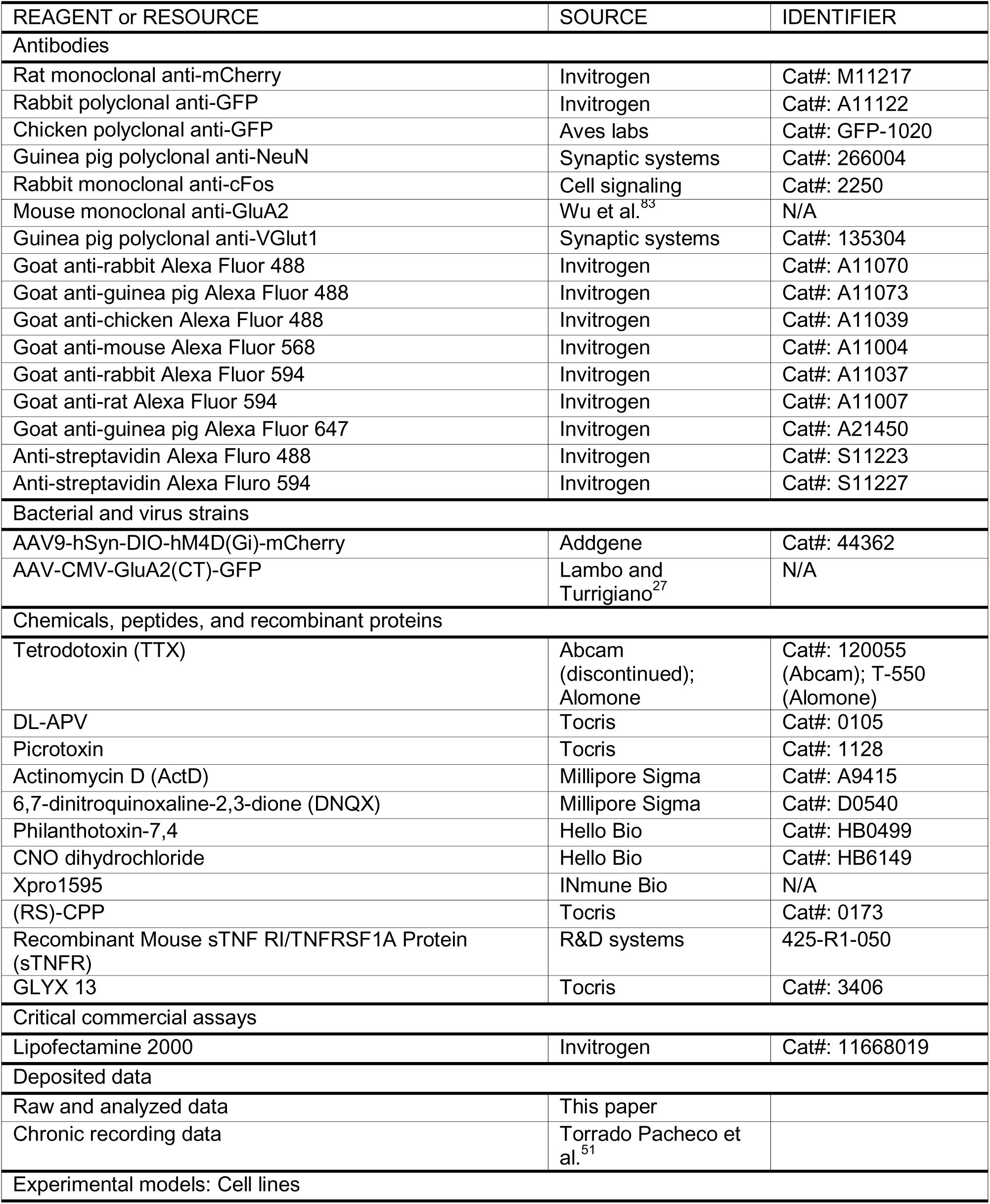
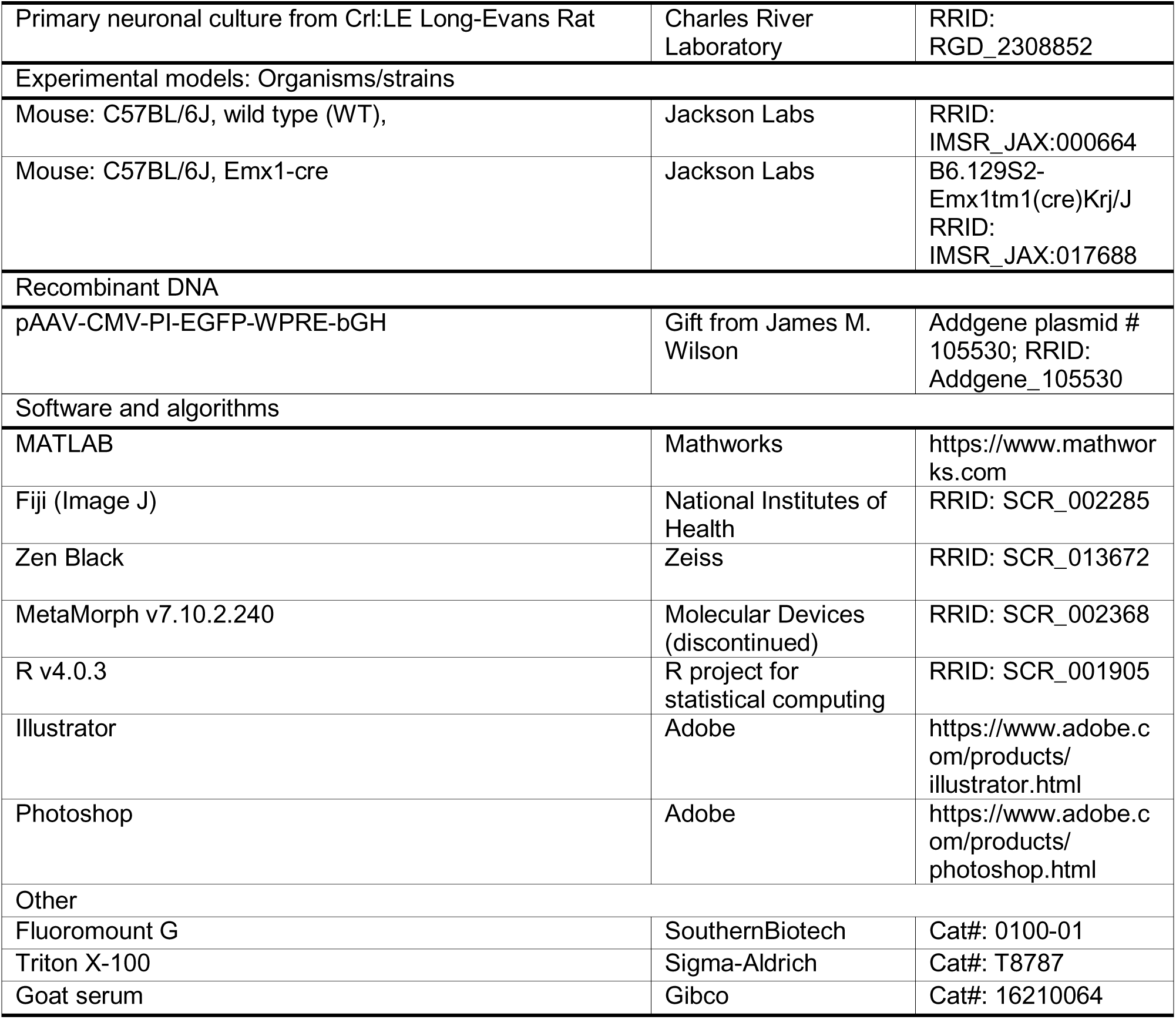

### Resource availability

#### Material availability

Materials used are listed in the Key Resource Table. No new unique reagent was generated.

#### Data and code availability

All data generated in this study are included in this article and can be accessed at Figshare: https://dx.doi.org/10.6084/m9.figshare.25883752. All scripts used for analysis have been deposited at the following URL: https://github.com/turrigianoCodeSpace/WeiWenManuscript2024.

### Experimental Model and Subject Details

All experimental procedures performed in this study strictly followed the protocols approved by the Brandeis Animal Care and Use Committee (IACUC) and Institutional Biosafety Committee (IBC), which complied with Guide for the Care and Use of Laboratory Animals from the National Institute of Health. Animals were housed on a 12:12 light/dark cycle with ad libitum access to food and water (except when experiments dictated otherwise, see below). Pups were weaned between postnatal days (P) 19-21 and housed with at least one littermate (except when experiments required single-housing). For all experiments, both males and females were used; no sex-dependent difference was observed and therefore results were not separated by sex. For in vitro primary neuronal cultures, newborn pups between P0-3 from timed-pregnant Long-Evans rats were used for dissociation. C57BL/6J mice were used for all ex vivo slice physiology experiments, which were performed between P24-32 to capture the classic rodent visual system critical period. The number of animals used for each experiment are given in the corresponding figure legend.

### Method Details

#### Primary neuronal culture

Primary neuronal cultures were dissociated from the visual cortex of newborn rat pups (P0-3) as previously described^8,83^. Briefly, pups were anesthetized with isofluorane and decapitated, and the visual cortex was removed and incubated at 37 °C with an enzyme solution containing 25 U/mL papain, 1 mM L-cystein, and 0.5 mM EDTA. The tissue was then rinsed and suspended with trypsin before being centrifuged at low speed for 5 minutes. The pellet was resuspended with neuronal medium supplemented with NS21^84^ and plated onto 35 mm glass-bottomed dishes pre-seeded with glial feeders. Cultures were incubated in a 5% CO_2_ incubator at 37 °C until being used for experiments, and were fed every 3-4 days by replacing 1 mL of old medium with fresh medium.

#### Drug treatment/administration

##### Cultured neurons

For electrophysiology experiments, primary neuronal cultures were treated with specific pharmacological agents between 9 and 14 days in vitro (DIV), usually up to 24 hours before being used for electrophysiology. Key drugs used in this study are listed in the Key Resource Table; all drugs were dissolved in Milli-Q water except for ActD, which was dissolved in 5% DMSO/PBS (working DMSO concentration 0.08%). The concentrations of drugs are as follows: TTX, 4 μM; DL-APV, 100 μM; Philanthotoxin, 10 μM; sTNFR, 2 μg/mL; GLYX-13, 1 μM; ActD, 50 μM.

For immunocytochemistry experiments, cultures were transfected with a plasmid encoding EGFP (500 ng per dish) using Lipofectamine 2000 (Invitrogen) on DIV8. After 48 hours, transfected cells were treated with DL-APV (100 μM) for 1 hour before being used for immunocytochemistry.

##### Animals

For DREADD activation, CNO dihydrochloride (CNO) was delivered to the animals via drinking water as described previously^29^. Briefly, CNO was dissolved in tap water to 0.05 mg/mL with 10 mM saccharine chloride added. Animals were given saccharine-only water with the same concentration for a day before being switched to a CNO-containing one. They were sacrificed for experiments after 24 hours of CNO administration unless indicated otherwise.

For other drug administrations including Xpro1595 and CPP, the drug was dissolved in 0.9% sterile saline to the desired concentration (Xpro1595, 1 mg/mL; CPP, 1.5 mg/mL). Appropriate amounts of the drug solution were then delivered to the animals via subcutaneous injection to reach desired dosages (Xpro1595, 10-20 mg/kg; CPP, 15 mg/kg).

#### Virus Injection surgery

Virus injection surgeries were performed between P14-19 on a stereotaxic apparatus while animals were anesthetized with ketamine/xylazine/acepromazine cocktail (KXA). The monocular region of the primary visual cortex (V1m) was targeted unilaterally using stereotaxic coordinates (Allen Brain Atlas) that were proportionally adjusted according to the age-dependent bregma-lambda distance difference. Unless noted otherwise, 200-300 nL of virus were delivered into the targeted area via a micropipette. Surgerized animals were allowed to recover in their home cages for a week before slice physiology experiments.

#### Transcardial perfusion

Animals were anesthetized with a triple dose of KXA used for virus injection surgery. They were then transported to a fume hood, and the heart was exposed, and a needle was inserted into the left ventricle. The right atrium was then cut open, and 1x PBS was perfused through the circulatory system using a peristaltic pump for 5 minutes. The PBS was switched to 4% paraformaldehyde (PFA) when the liver color turned into light-brown, and the perfusion continued for about 5 minutes. The brain was then removed and incubated in 4% PFA with shaking at 4C overnight.

#### Immunostaining

For acute slices harvested from electrophysiology experiments, following slice physiology recordings, slices were post-fixed in 4% (w/v) PFA overnight and transferred to 1x PBS for storage before staining. For the cFos experiment in Figure S6C, the fixed brain was mounted on a Leica VT1000S vibratome (Leica Biosystems, Buffalo Grove IL), and V1 was sectioned into 60 µm slices. Slices were washed with 1x PBS for at least 30 min (6 x 5 min) before being incubated in a blocking solution (0.1% Triton X-100, 0.05% NaN_3_, and 1% BSA in 1x PBS) for 1 hour. Blocked slices were then incubated in the same blocking solution at 4 °C for 24 hours with the following primary antibodies (1:1000), respectively: rabbit anti-GFP (if applicable) and rat anti-mCherry for acute slices, rabbit anti-cFos and guniea pig anti-NeuN for the cFos experiment. The following day, slices were washed for 30 min in PBS and then incubated in a solution (0.05% NaN_3_ and 1% BSA in 1x PBS) containing the following secondary antibodies (1:500), respectively: goat anti-rabbit Alexa Fluor 488 (if applicable), anti-rat Alexa Fluor 594, and anti-streptavidin Alexa Fluro 488 or 594 for acute slices, goat anti-rabbit Alexa Fluro 594 and goat anti-guinea pig Alexa Fluor 488 for the cFos experiment. Acute slices were incubated at 4°C overnight, all other slices were incubated at room temperature for 3 hours. Finally, slices were washed for another 30 min before mounted in Fluoromount-G mounting medium.

For cultured neurons, 48 hours post-transfection, cultured neurons were fixed with 4% (w/v) PFA with 5% sucrose for 10 minutes, then washed with 1x PBS (3 x 5 min). Next, neurons were incubated in blocking solution (5% goat serum in 1x PBS) for 30 min to prevent non-specific binding. To visualize surface GluA2 (sGluA2) expression, neurons were incubated with mouse anti-GluA2 (1:500) in blocking solution for 90 min under non-permeabilized conditions, as described previously^8,83^. Afterward, neurons were washed with 1x PBS (3 x 5 min). The subsequent steps were performed under permeabilized conditions. Neurons were incubated in permeabilization buffer (0.2% Triton X-100 in 1x PBS) for 15 mins followed by a second blocking step for an additional 30 mins. Neurons were then incubated overnight at 4°C in antibody dilution buffer (0.1% Triton X-100 and 5% goat serum in 1x PBS) with the following primary antibodies: chicken anti-GFP and guinea pig anti-VGlut1 (1:1000). The next day, neurons were washed three times in 1x PBS for 5 min each, then incubated for 1 hour in antibody dilution buffer containing the following Alexa-Fluor-conjugated secondary antibodies: goat anti-chicken Alexa-488, goat anti-mouse Alexa-568, and goat anti-guinea pig Alexa-647 (1:400). After three more 5 min washes with 1x PBS, the glass coverslips containing immunostained neurons were detached from their dishes, mounted onto slides with Fluoromount-G medium, and sealed with nail polish.

Immunostained slices were imaged using an inverted Zeiss LSM 880 laser scanning confocal microscope using a x20 air objective. V1 and patched neurons (if applicable) were located based on the white matter morphology, and z-stacked images were acquired. Immunostained cultured neurons were imaged using an inverted Zeiss LSM880 Airyscan laser scanning confocal microscope, using a 63× oil immersion objective. Pyramidal neurons were identified during image acquisition based on characteristic morphological features, including a conical-shaped soma and distinct apical and basal dendrites. Z-stacked images of dendritic branches from the apical dendrites were acquired and processed with Airyscan. Maximum intensity projections were generated using ZEN Black software.

#### Ex vivo acute slice preparation

Animals were anesthetized with isoflurane. After toe-pinch check, the animal was decapitated and coronal slices (300 µm) containing V1m from both hemispheres were obtained using a Leica VT1000S vibratome. Slices were first transferred to an oxygenated chamber filled with choline solution (in mM: 110 Choline-Cl, 25 NaHCO_3_, 11.6 Na-Ascorbate, 7 MgCl_2_, 3.1 Na-Pyruvate, 2.5 KCl, 1.25NaH_2_PO_4_, and 0.5 CaCl_2_, osmolarity adjusted to 310 mOsm with dextrose, pH 7.4) for recovery, and then transferred back to oxygenated standard artificial cerebrospinal fluid (ACSF, in mM: 126 NaCl, 25 NaHCO_3_, 3 KCl, 2 CaCl_2_, 2 MgSO_4_, 1 NaH_2_PO_4_, 0.5 Na-Ascorbate, osmolarity adjusted to 310 mOsm with dextrose, pH 7.4) and incubated for 40 min. Slices were used for electrophysiology 1-5 hours post slicing.

#### Electrophysiology

##### Experimental setup

For culture electrophysiology, 35 mm glass-bottom dishes with primary neuronal culture were rinsed with ACSF (in mM: 126 NaCl, 25 NaHCO_3_, 3 KCl, 2 CaCl_2_, 2 MgSO_4_, 1 NaH_2_PO_4_, 0.5 Na-Ascorbate, osmolarity adjusted to 320 mOsm with dextrose, pH 7.4) and placed on an Olympus IX70 upright fluorescence microscope. Neurons were superfused in oxygenated ACSF at 32° C throughout the experiment. Pyramidal neurons were visually identified by their morphology under a 20x objective, and then approached and patched under a 40x objective. Borosilicate glass pipettes with resistance between 3-7 MΩ were filled with KMeSO_4_-based internal solution (in mM: 120 KMeSO_4_, 10 KCl, 10 HEPES, 0.5 EGTA, 2 MgSO_4_, 10 Na-Phosphocreatine, 3 K_2_-ATP, and 0.3 Na-GTP, osmolarity adjusted to 310 mOsm with dextrose, pH adjusted to 7.4 with KOH).

For slice electrophysiology, coronal slices containing V1 were placed on an Olympus BX51WI upright epifluorescence microscope equipped with infrared-DIC optics. V1m was identified as previously described^29^. Pyramidal neurons were visually targeted for whole-cell recordings using a 40x water-immersion objective; visual identification was based on the teardrop shaped somata and the presence of an apical dendrite, and morphology was confirmed post hoc from biocytin fill reconstructions. Neurons expressing exogeneous proteins were identified by the fluorescent marker embedded in the corresponding constructs. Borosilicate glass pipettes with resistance between 4 to 6 MΩ were filled with K^+^ Gluconate-based internal solution (in mM: 100 K-gluconate, 10 KCl, 10 HEPES, 5.37 Biocytin, 0.5 EGTA, 10 Na-Phosphocreatine, 4 Mg-ATP, and 0.3 Na-GTP, osmolarity adjusted to 295 mOsm with sucrose, pH adjusted to 7.4 with KOH). All recordings were performed in slices that were superfused in oxygenated standard ACSF at 34 °C.

All electrophysiological recordings were performed using a Multiclamp 700B amplifier with a CV-7B headstage (Molecular Devices, Sunnyvale CA). Data were passed through a 6 kHz Bessel low-pass filter and acquired at 10 kHz using a National Instruments Data Acquisition Board (DAQ, National Instruments, Woburn, MA) and an open-source MATLAB-based software WaveSurfer (HHMI Janelia, Ashburn VA).

##### mEPSC recordings

For spontaneous mEPSC recordings, cultured neurons or slices were superfused with standard ASCF containing a drug cocktail of tetrodotoxin (TTX, 0.1 µM), D-2-amino-5-phosphonovalerate (APV, 50 µM), and picrotoxin (25 µM) to isolate mEPSCs. Pyramidal neurons were targeted and held at −70 mV in whole-cell voltage clamp, and series resistance was not compensated. Each neuron was recorded for 3-5 minutes in a series of 30s segments, and a 500 ms 5 mV hyperpolarizing voltage step was applied at the beginning of each segment so that passive properties could be monitored throughout the recording. Neurons were excluded if series resistance was > 20 MΩ, input resistance was < 100 MΩ (70 MΩ for culture), membrane potential was > −55 mV at break-in (−50 mV for culture), or these properties changed by > 15%.

##### Intrinsic excitability measurements

For intrinsic excitability measurements, cultured neurons or slices were superfused with standard ACSF containing APV (50 µM), picrotoxin (25 µM), and DNQX (25 µM) to block synaptic currents. Pyramidal neurons were held in current clamp with a small DC current injection to maintain the resting membrane potential at −60 mV (culture) or −70 mV (slice). Pipette capacitance was then neutralized, and bridge balance was compensated. Frequency versus current (F-I) curves were obtained by delivering a series of 0.5 s (culture) or 1 s (slice) long current injections in amplitude increments of 25 pA (culture) or 20 pA (slice). To monitor passive neuronal properties (except the resting membrane potential) throughout the recording, a 500ms 50 pA hyperpolarizing current step was applied before each current injection step. To monitor resting membrane potential changes, neurons were held in current clamp without any current injection and recorded for 1 minute before and after F-I curve measurements. To measure rheobase currents for cultured neurons, the rheobase was first estimated by locating the current step that had evoked the first spike during the F-I curve measurement, then 5 0.5 s long current steps starting from the previous lower current step and with amplitude increments of 5 pA were injected until a spike was evoked. Neurons that did not meet the criteria listed above for mEPSC recordings were excluded; in addition, neurons were excluded if the deviation of baseline membrane potential from their corresponding holding potentials was > 5% or the voltage firing threshold was >-25 mV.

##### Spontaneous firing recordings

To measure spontaneous firing in dissociated cultures, cultured neurons were superfused with either plain ACSF or with 50 µM APV. Cells were approached and broken into whole-cell mode under voltage clamp, and recordings were obtained under current clamp where a small DC current was injected to maintain the membrane potential around −50 mV. Each cell was recorded for 3-5 minutes in a series of 30s segments, and a 500 ms 50 pA hyperpolarizing voltage step was applied at the beginning of each segment so that passive properties could be monitored throughout the recording. Neurons were excluded if access resistance was > 20 MΩ, input resistance was < 70 MΩ, resting membrane potential was > −50 mV at break-in, or any property during the recording changed by > 15%.

### Quantification and statistical analysis

All data obtained in this study were analyzed using in-house scripts written in MATLAB except the immunohistochemistry images, which were analyzed using Image J. Results illustrated via box plots are reported as median with 25^th^ and 75^th^ percentiles, with individual data points shown on the side; results shown in other types of plot are reported as described in the corresponding figure legends. Effect sizes are reported as a percentage of the group of interest (indicated in either the main text or the legend); samples sizes (the number of animals or dissociations used for each experiment and the number of neurons collected for each condition), statistical tests performed, and p values are given in either the corresponding results section or the figure legends.

#### mEPSC recordings

Spontaneous mEPSC events were automatically detected using an in-house MATLAB script (modified from Miska et al.^85^). Briefly, the script filters the raw recording trace and then slides a mEPSC event-shaped template to find regions that fit the detection criteria^86^. Detected putative mEPSC events were then passed through multiple quality control modules to exclude the ones that were determined to be false-positive. For culture recordings, rise time cutoff was set to 2 ms; for slice recordings, rise time cutoff for detection was lowered to 1 ms to minimize the inclusion of highly filtered events from distal dendrites. Mean amplitude and frequency were first calculated for each 30 s of recording, which were then averaged to give the mean value for each neuron. Rise time and decay time constants were calculated from the waveform average traces for each neuron. Rise time is defined as the time for the current to increase from 10% to 90% of the peak amplitude. Decay time constant (τ) is derived from a first-order exponential fit of the decay phase. To generate the cumulative distribution and the average waveform for each condition, 100 events were randomly selected from each neuron and pooled.

#### Spontaneous firing recordings

Raw spontaneous firing recordings were used for analysis without any filtering. Spikes were automatically detected using an in-house MATLAB script mainly based on the maximum dV/dt (>20 V/s) and the peak amplitude (> 0 mV). For each neuron, the mean firing rates were calculated for each 30s trace, which were then averaged to get the mean firing rate of the cell. Neurons were excluded if the mean spontaneous firing rate was < 0.01 Hz.

#### Intrinsic excitability recordings

Raw recordings obtained during the intrinsic excitability measurement were used for analysis without any filtering. For traces with depolarizing current injections, spikes were automatically detected using an in-house MATLAB script mainly based on the maximum dV/dt (> 20 V/s) and the peak amplitude (> 0 mV). Definitions of single spike or spike train properties analyzed in this study were primarily adapted from the electrophysiology technical white paper published by Allen Institute (https://celltypes.brain-map.org).

Briefly, for a given current step with a spike train, instantaneous firing rate (IFR) is defined as the mean reciprocal of the first two inter-spike intervals; mean instantaneous firing rate (mean IFR) is defined as the mean reciprocal of all inter-spike intervals; latency is defined as the time difference between the current step onset and the first spike; spike frequency adaptation is defined as the rate at which firings speeds up or slows down during the current step, and calculated as the mean normalized difference between two consecutive inter-spike intervals (higher values indicate more adaptation). For a given neuron, rheobase is defined as the smallest depolarizing current injection that has evoked one or more spikes.

For single spike properties, the first spike evoked at the rheobase is used for analysis unless noted otherwise. Spike threshold is defined as the membrane potential where dV/dt exceeds 5% of the maximum dV/dt during the depolarizing phase (Wen and Turrigiano^29^). Spike width at half-maximum is defined as the full width at half maximum of the spike height. Afterhyperpolarization (AHP) amplitude is defined as the difference between the minimum membrane potential reached after a spike and the mean membrane potential between the current step onset and the spike threshold (normalized to the spike height of the same cell).

#### Image analysis

To quantify the number of cFos-positive neurons in mouse V1, images obtained from post-staining slices (against cFos and NeuN) were first background-subtracted and thresholded against the mean intensity of the NeuN signal from 3 negative control slices imaged in the same session (secondary antibodies only). A region of interest (ROI) of 250×200 μm was then manually selected from V1, and cFos signal was automatically background-subtracted. Cell somas were outlined in the NeuN channel, and cFos signals were measured in each identified cell soma. Cells that showed non-zero mean intensity values in the cFos channel were considered cFos-positive neurons. For each animal, 10-12 ROIs that spanned V1 were selected, and the percentage of cFos-positive neurons was reported for each ROI.

To perform synaptic puncta colocalization and intensity quantification in cultured neurons, images were processed using the MetaMorph software (Molecular Devices) as previously described^8,83^. A region of interest was manually drawn around the dendritic branches, and the granularity function in MetaMorph was applied to threshold, mask, and binarize the synaptic puncta. Integrated granule intensity, representing the sum of pixel intensities across puncta, was used to quantify synaptic sGluA2 intensity. Synapses were defined as puncta labeled with both sGluA2 and VGlut1.

#### Pairwise correlation analysis

The data set used for correlation analysis was obtained from chronic multielectrode recordings from the control hemispheres of V1m of freely behaving Long-Evans rats, between the ages of P21 and P36, obtained in a previous study^51^. Only animals that had at least two regular spiking units (RSUs) were included in the analysis. In total, 17 regular spiking units (RSUs) from 5 animals were used. Pairwise correlation coefficients of V1 neurons were calculated as previously described^15,54^. Briefly, the spike timestamps for each neuron were divided into 100 ms bins, and the spike count was computed for each bin, which generated a spike count vector for each neuron. For each animal, the Pearson correlation coefficient r of spike counts was computed for a neuronal pair in 30-minute episodes with a 5-minute sliding window, producing 139 values for each pair in a 12-hour session. These values were then normalized to the mean correlation value of a selected baseline window (see Figure 7A legend). Example correlation matrices were generated from 5 neurons from 1 animal.

#### Statistical analysis

All statistical tests and p values for each dataset are given in the figure legends. Statistical tests were determined by the experimental design and planned comparisons. For most of our experiments using neuronal culture (i.e., Figures 1E, 4C, 4G, 6B, 6E, S1A-F, S2C-F, S4B-H, and S5A-I), we performed multiple drug treatments and recordings in parallel using sister cultures, and planned comparisons between multiple groups. Therefore, we chose a multivariant test (Kruskal-Wallis with Tukey post hoc correction) for these datasets. To determine the impact of Xpro (Figure 2) or CPP (Figure 5) on synaptic scaling or IHP we used a within-animal design, where we planned comparisons from activity-deprived and unmanipulated V1m hemispheres from the same animals (data in Figures 2C, 2I, 2J, 5D-G, S2G-I, S3B-F, and S3H); for these datasets we used two-way ANOVA followed by Tukey post hoc correction to test for interaction between two factors (CNO vs. DR+CNO and Saline vs. Xpro/CPP) after confirming they passed an Anderson-Darling normality test. Additionally, for cellular properties that did not pass the normality test (Figures S3G and S3I), we performed a 4-way multivariant test (Kruskal-Wallis with Tukey post hoc correction). To differentiate L/D and circadian effects on IHP we planned a 3-way comparison (Figures 7D, S6F, S6H-I, and S7A-D), using littermate animal, and similarly we planned a 3-way comparison to determine if GluA2 C-tail expression affects IHP (Figures 3E-H); therefore we chose a multivariant test (Kruskal-Wallis with Tukey post hoc correction) for these datasets. To determine the impact of CPP on Light-induced IHP against vehicle control (Figures 7G-I and S7E-H), the pairwise correlation values across light and dark (Figure S6B), the impact of CPP on cFos activity (Figure S6D), and the impact of APV on synaptic AMPAR accumulation (Figure S4J), we used a pairwise test. For unplanned post hoc comparisons, such as the impact of treatments on cell capacitance, we performed a multivariant test corrected for multiple comparisons (Figure S5J). For distribution comparisons (Figures 2D-E, S6G, and S6K), a two-sample Kolmogorov-Smirnov test was used. Reported P values were rounded to the nearest hundredth, or for very small values were reported in scientific notation with 2 significant figures. Results were considered significant if p <= 0.05.

## Supplemental Information

Figures S1-S7. Table S1.

