## Supplementary figures 1-7; supplementary table 1 for "Modular Arrangement of Synaptic and Intrinsic Homeostatic Plasticity within Visual Cortical Circuits"

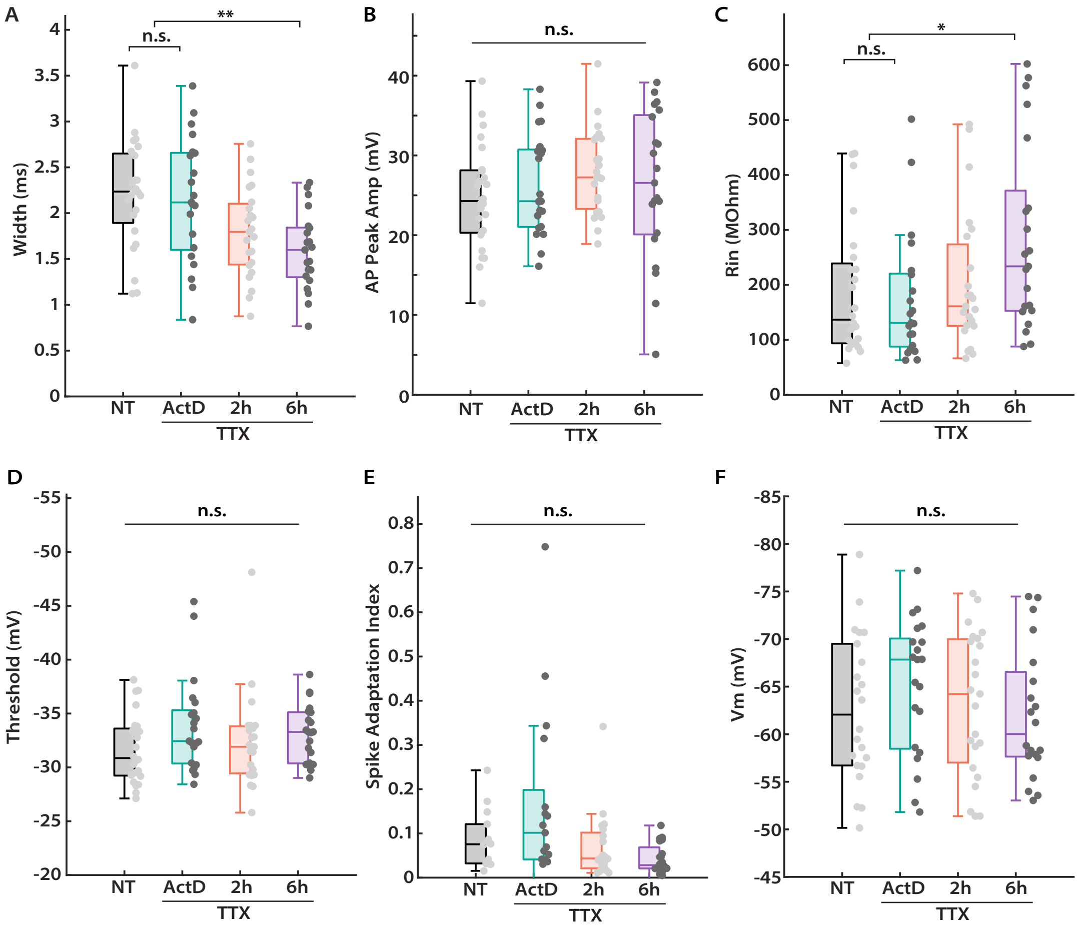
A
B
C
D
E
F
Width (ms)
AP Peak Amp (mV)
Rin (MOhm)
Threshold (mV)
Spike Adaptation Index
Vm (mV)
n.s.
**
*
NT
ActD
2h
6h
TTX

Figure S1. Cellular properties across the time course of IHP expression in cultured neocortical pyramidal neurons. Related to Figure 1.

(A) Comparison of full width at half-maximum for the first spike at rheobase from the indicated conditions. Kruskal-Wallis test with Tukey correction: NT vs. ActD, $p = 0.97$; NT vs. 2h, $p = 0.08$; NT vs. 6h, $p = 2.7E-3$; ActD vs. 2h, $p = 0.25$; ActD vs. 6h, $p = 0.02$; 2h vs. 6h, $p = 0.65$.

(B) Comparison of the peak amplitude for the first spike at rheobase from the indicated conditions. Kruskal-Wallis test: $p = 0.45$.

(C) Comparison of the neuronal input resistance from the indicated conditions. Kruskal-Wallis test with Tukey correction: NT vs. ActD, $p = 0.98$; NT vs. 2h, $p = 0.92$; NT vs. 6h, $p = 0.02$; ActD vs. 2h, $p = 0.74$; ActD vs. 6h, $p = 0.01$; 2h vs. 6h, $p = 0.33$.

(D) Comparison of voltage spiking threshold for the first spike at rheobase from the indicated conditions. Kruskal-Wallis test: $p = 0.17$.

(E) Comparison of spike frequency adaptation at 175 pA of current injection from the indicated conditions. Kruskal-Wallis test: $p = 0.10$.

(F) Comparison of resting membrane potential from the indicated conditions. Kruskal-Wallis test: $p = 0.14$.

A through F: same sample sizes as shown in Figure 1E.

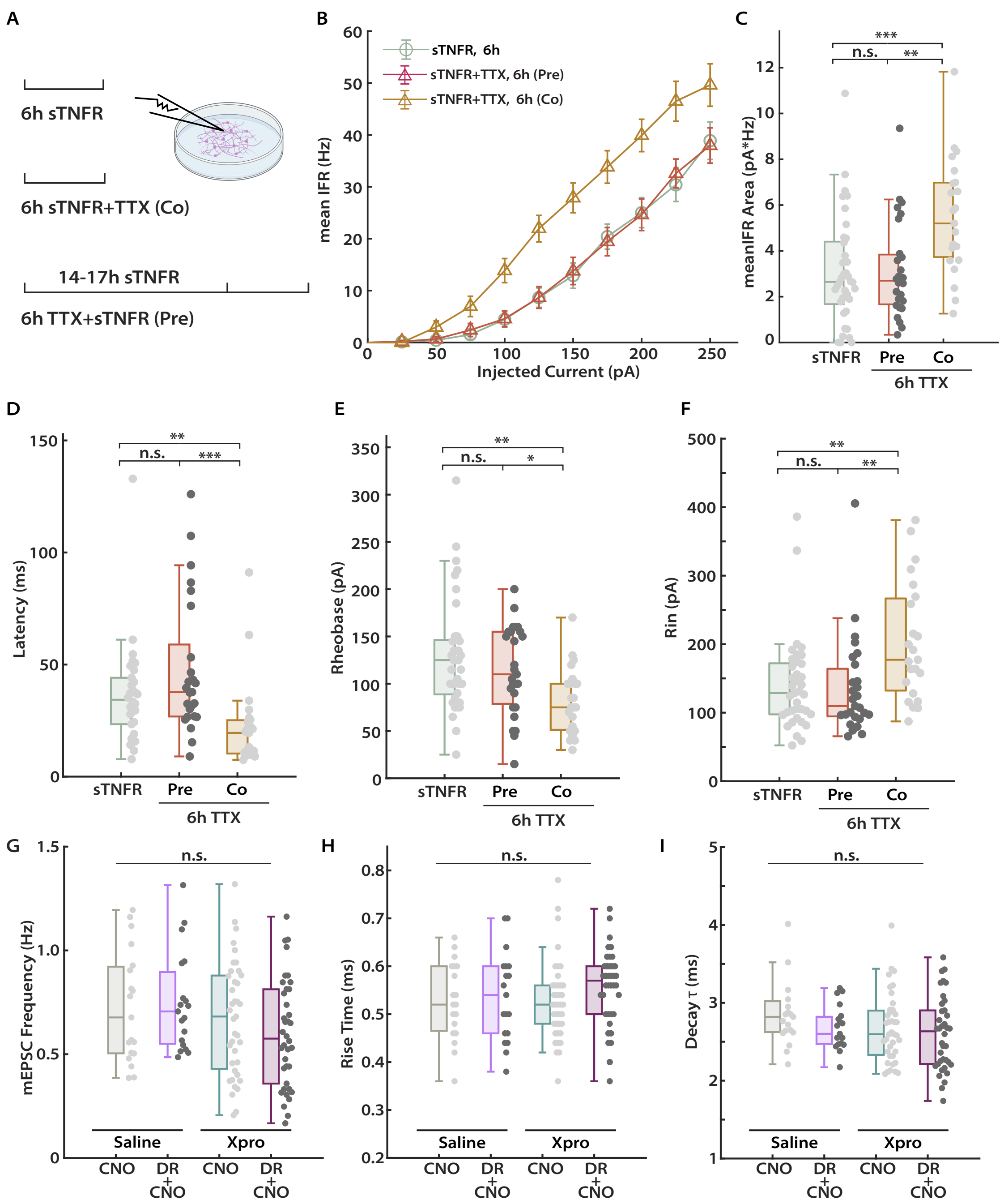
A
6h sTNFR
6h sTNFR+TTX (Co)
14-17h sTNFR
6h TTX+sTNFR (Pre)
B
sTNFR, 6h
sTNFR+TTX, 6h (Pre)
sTNFR+TTX, 6h (Co)
mean IFR (Hz)
Injected Current (pA)
C
meanIFR Area (pA*Hz)
n.s.
**
***
sTNFR
Pre
Co
6h TTX
D
Latency (ms)
E
Rheobase (pA)
*
F
Rin (pA)
G
mEPSC Frequency (Hz)
Saline
Xpro
CNO
DR + CNO
H
Rise Time (ms)
I
Decay τ (ms)

Figure S2. IHP expression *in vitro* requires TNF$\alpha$ signaling. Related to Figure 2.

(A) Experimental paradigm. Cultured neocortical neurons were treated with sTNFR alone, or together with TTX (Co), or pretreated with sTNFR for 14-17 hours before adding TTX (Pre).

(B) Comparison of F-I curves for pyramidal neurons from the indicated conditions in A. sTNFR, n = 37, 6 dissociations; Pre, n = 27, 3 dissociations; Co, n = 23, 3 dissociations.

(C) Comparison of the area under F-I curve for each neuron from the indicated conditions. Kruskal-Wallis test with Tukey correction: sTNFR vs. Pre, p = 0.99; sTNFR vs. Co, p = 7.9E-4; Pre vs. Co, p = 2.6E-3.

(D) Comparison of latency to the first spike at 175 pA of current injection from the indicated conditions. Kruskal-Wallis test with Tukey correction: sTNFR vs. Pre, p = 0.50; sTNFR vs. Co, p = 3.0E-3; Pre vs. Co, p = 8.0E-5.

(E) Comparison of rheobase current from the indicated conditions. Kruskal-Wallis test with Tukey correction: sTNFR vs. Pre, p = 0.84; sTNFR vs. Co, p = 2.4E-3; Pre vs. Co, p = 0.02.

(F) Comparison of neuronal input resistance from the indicated conditions. Kruskal-Wallis test with Tukey correction: sTNFR vs. Pre, p = 0.86; sTNFR vs. Co, p = 7.4E-3; Pre vs. Co, p = 3.0E-3.

(G) Comparison of the mean mEPSC frequency for L2/3 pyramidal neurons from the indicated conditions. Two-way ANOVA test: interaction, p = 0.36; CNO vs. DR+CNO on all groups, p = 0.62; Saline vs. Xpro on all groups: p = 0.06.

(H) Comparison of the mEPSC rise time for L2/3 pyramidal neurons from the indicated conditions. Two-way ANOVA test: interaction, p = 0.54; CNO vs. DR+CNO on all groups, p = 0.26; Saline vs. Xpro on all groups: p = 0.55.

(I) Comparison of the mEPSC decay time constant ($\tau$) for L2/3 pyramidal neurons from the indicated conditions. Two-way ANOVA test: interaction, p = 0.17; CNO vs. DR+CNO on all groups, p = 0.21; Saline vs. Xpro on all groups: p = 0.14.

G through I: same samples sizes as shown in Figure 2C.

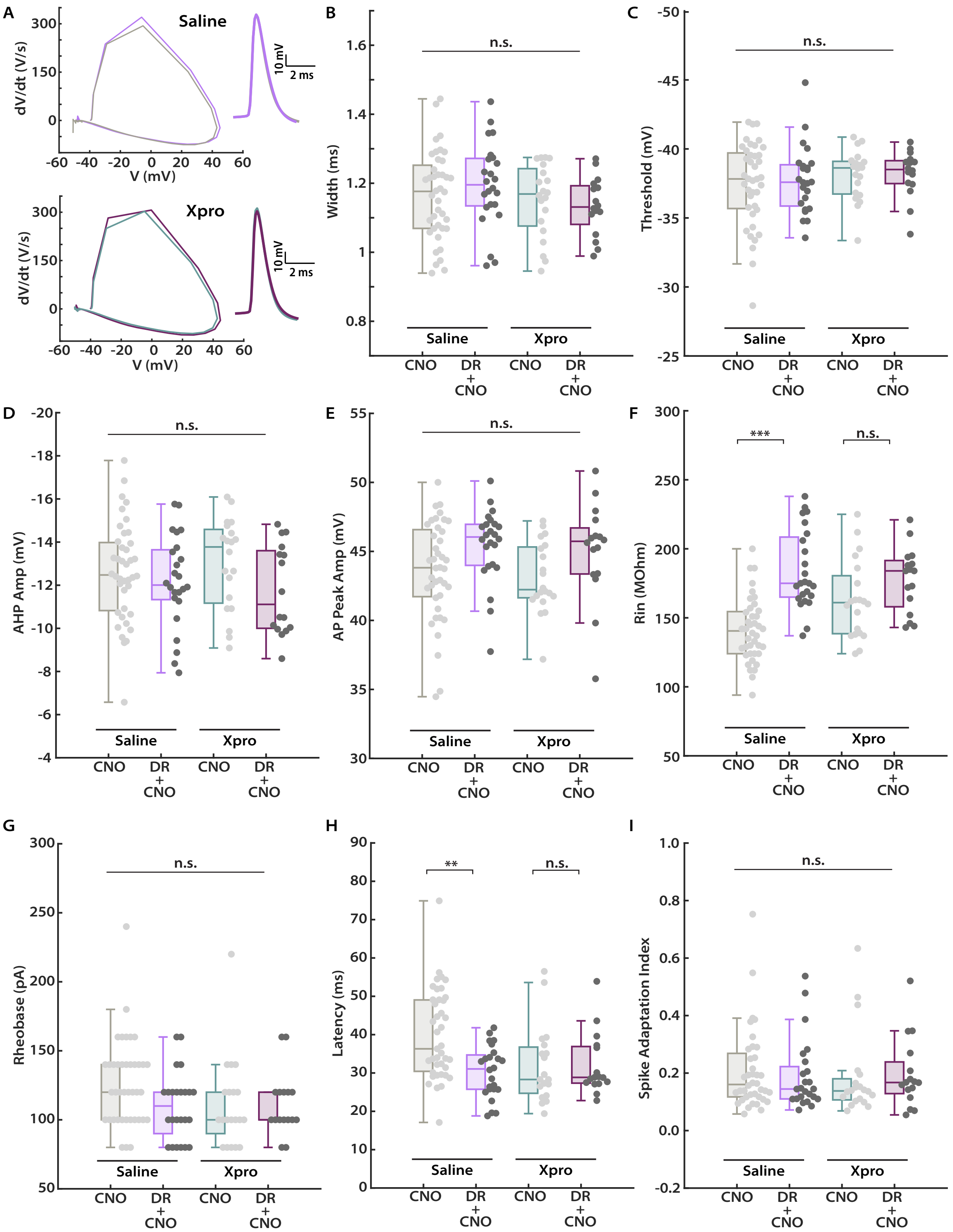
A
Saline
Xpro
dV/dt (V/s)
V (mV)
10 mV
2 ms
B
Width (ms)
C
Threshold (mV)
D
AHP Amp (mV)
E
AP Peak Amp (mV)
F
Rin (MOhm)
G
Rheobase (pA)
H
Latency (ms)
I
Spike Adaptation Index
n.s.
***
**
CNO
DR + CNO

Figure S3. Cellular and spike parameters following IHP induction in V1 L2/3 pyramidal neurons. Related to Figure 2.

(A) Phase plane plots (left, action potential plotted as dV/dt vs. membrane potential) for the average waveforms (right, overlay of both CNO and DR+CNO conditions) of the first spike evoked at rheobase from animals treated with saline (upper) and Xpro (lower), respectively.

(B) Comparison of full width at half-maximum for the first spike at rheobase from the indicated conditions. Two-way ANOVA test: interaction, p = 0.33; CNO vs. DR+CNO on all groups, p = 0.78; Saline vs. Xpro on all groups: p = 0.12.

(C) Comparison of action potential voltage threshold for the first spike at rheobase from the indicated conditions. Two-way ANOVA test: interaction, p = 0.86; CNO vs. DR+CNO on all groups, p = 0.71; Saline vs. Xpro on all groups: p = 0.30.

(D) Comparison of afterhyperpolarization amplitude for the first spike at rheobase from the indicated conditions. Two-way ANOVA test: interaction, p = 0.39; CNO vs. DR+CNO on all groups, p = 0.07; Saline vs. Xpro on all groups: p = 0.99.

(E) Comparison of spike peak amplitude for the first spike at rheobase from the indicated conditions. Two-way ANOVA test: interaction, p = 0.96; CNO vs. DR+CNO on all groups, p = 5.7E-3; Saline vs. Xpro on all groups: p = 0.49. Tukey correction: CNO+Saline vs. DR+CNO+Saline, p =0.11; CNO+Saline vs. CNO+Xpro, p = 0.94; CNO+Saline vs. DR+CNO+Xpro, p = 0.42; DR+CNO+Saline vs. CNO+Xpro, p = 0.07; DR+CNO+Saline vs. DR+CNO+Xpro, p = 0.97; CNO+Xpro vs. DR+CNO+Xpro, p = 0.27.

(F) Comparison of neuronal input resistance from the indicated conditions. Two-way ANOVA test: interaction, p = 9.8E-3; CNO vs. DR+CNO on all groups, p = 0.16; Saline vs. Xpro on all groups: p = 0.05. Tukey correction: CNO+Saline vs. DR+CNO+Saline, p =1.5E-8; CNO+Saline vs. CNO+Xpro, p = 0.01; CNO+Saline vs. DR+CNO+Xpro, p = 2.6E-5; DR+CNO+Saline vs. CNO+Xpro, p = 0.03; DR+CNO+Saline vs. DR+CNO+Xpro, p = 0.85; CNO+Xpro vs. DR+CNO+Xpro, p = 0.31.

(G) Comparison of rheobase current from the indicated conditions. Kruskal-Wallis test with Tukey correction: CNO+Saline vs. DR+CNO+Saline, p =0.14; CNO+Saline vs. CNO+CPP, p = 0.10; CNO+Saline vs. DR+CNO+CPP, p = 0.29; DR+CNO+Saline vs. CNO+CPP, p = 0.99; DR+CNO+Saline vs. DR+CNO+CPP, p = 1.0; CNO+CPP vs. DR+CNO+CPP, p = 0.99.

(H) Comparison of latency to the first spike at 200 pA of current injection from the indicated conditions. Two-way ANOVA test: interaction, p = 0.02; CNO vs. DR+CNO on all groups, p = 0.02; Saline vs. Xpro on all groups: p = 0.15. Tukey correction: CNO+Saline vs. DR+CNO+Saline, p = 1.6E-3; CNO+Saline vs. CNO+Xpro, p = 0.03; CNO+Saline vs. DR+CNO+Xpro, p = 0.04; DR+CNO+Saline vs. CNO+Xpro, p = 0.93; DR+CNO+Saline vs. DR+CNO+Xpro, p = 0.94; CNO+Xpro vs. DR+CNO+Xpro, p = 1.0.

(I) Comparison of spike frequency adaptation at 400 pA of current injection from the indicated conditions. Kruskal-Wallis test: p = 0.75.

A through I: same sample sizes as shown in Figures 2G and 2I.

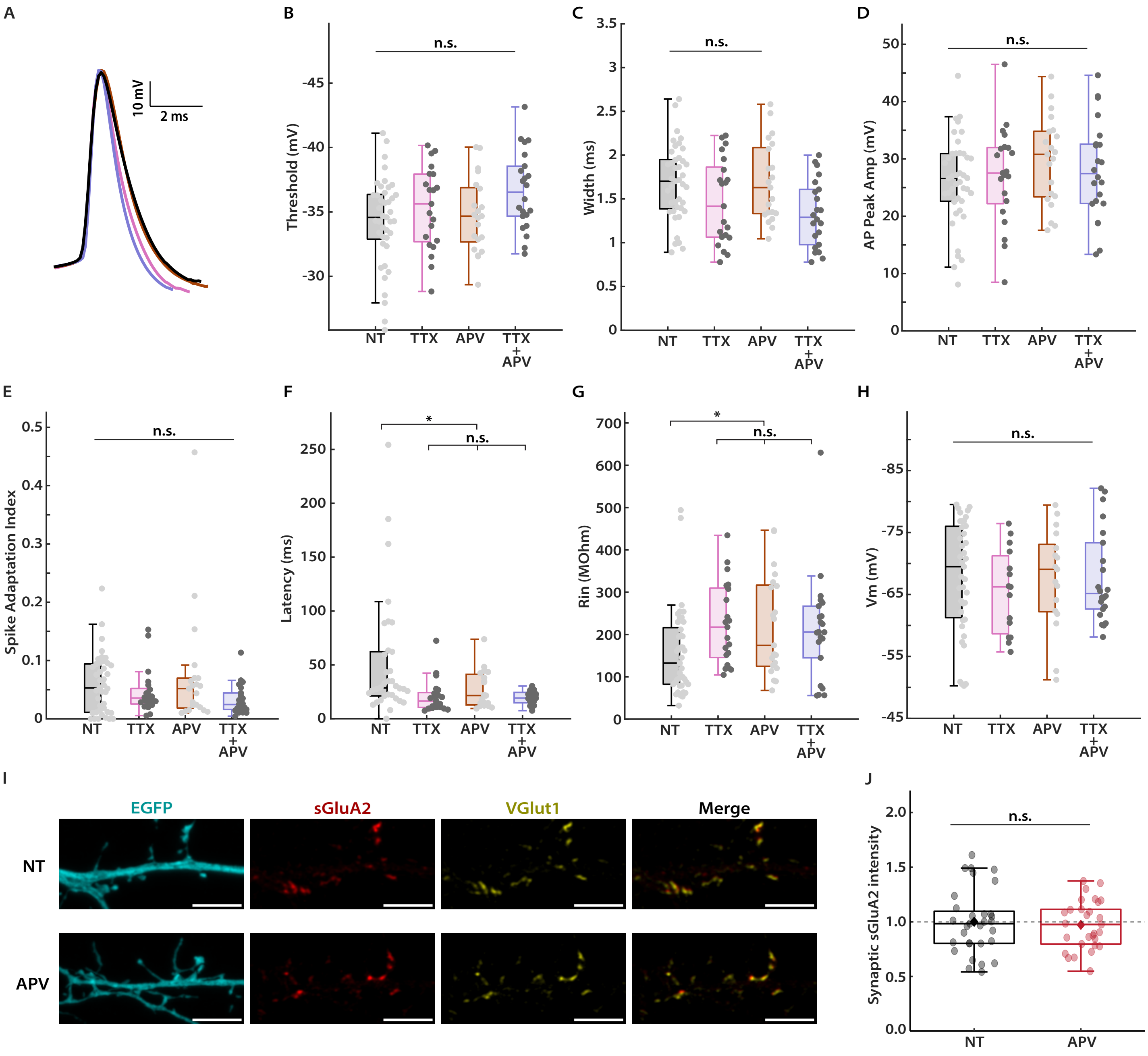
A
10 mV
2 ms
B
n.s.
Threshold (mV)
NT
TTX
APV
TTX + APV
C
Width (ms)
D
AP Peak Amp (mV)
E
Spike Adaptation Index
F
*
Latency (ms)
G
Rin (MOhm)
H
Vm (mV)
I
EGFP
sGluA2
VGlut1
Merge
NT
APV
J
Synaptic sGluA2 intensity

Figure S4. Cellular properties that underlie IHP expression *in vitro*. Related to Figure 4.

(A) Overlay of peak-scaled average waveforms of first spikes evoked at the rheobase from the indicated conditions (same color code as in Figure 4B)

(B) Comparison of action potential voltage threshold for the first spike at rheobase from the indicated conditions. Kruskal-Wallis test: p = 0.09.

(C) Comparison of full width at half-maximum for the first spike at rheobase from the indicated conditions. Kruskal-Wallis test with Tukey correction: NT vs. TTX, p = 0.29; NT vs. APV, p = 1.0; NT vs. TTX+APV, p = 0.01; TTX vs. APV, p = 0.52; TTX vs. TTX+APV, p = 0.70; APV vs. TTX+APV, p = 0.07.

(D) Comparison of the peak amplitude for the first spike at rheobase from the indicated conditions. Kruskal-Wallis test: p = 0.47.

(E) Comparison of spike frequency adaptation at 250 pA of current injection from the indicated conditions. Kruskal-Wallis test: p = 0.06.

(F) Comparison of latency to the first spike at 150 pA of current injection from the indicated conditions. Kruskal-Wallis test with Tukey correction: NT vs. TTX, p = 1.0E-3; NT vs. APV, p = 0.05; NT vs. TTX+APV, p = 0.01; TTX vs. APV, p = 0.46; TTX vs. TTX+APV, p = 0.97; APV vs. TTX+APV, p = 0.76.

(G) Comparison of neuronal input resistance from the indicated conditions. Kruskal-Wallis test with Tukey correction: NT vs. TTX, p = 0.013; NT vs. APV, p = 0.03; NT vs. TTX+APV, p = 0.06; TTX vs. APV, p = 0.98; TTX vs. TTX+APV, p = 0.90; APV vs. TTX+APV, p = 0.99.

(H) Comparison of resting membrane potential from the indicated conditions. Kruskal-Wallis test: p = 0.71.

(I) Representative images of apical dendrites from rat pyramidal neurons transfected with EGFP (cyan), comparing postsynaptic surface expression of GluA2 (sGluA2, red) and presynaptic expression of VGlut1 (yellow) in untreated (NT) and APV-treated (60 min) neurons. Scale bar, 5 μm.

(J) Quantification of synaptic sGluA2 intensity for conditions described in (I). Intensity values are normalized to the average sGluA2 intensity of untreated (NT) neurons for each dissociation. Each data point indicates the average synaptic sGluA2 intensity for an individual neuron. The diamond marker indicates the average of three separate dissociations. NT, n = 31; APV (60 min): n = 29. Unpaired t-test: p = 0.68.

A through H: same sample sizes as shown in Figure 4C.

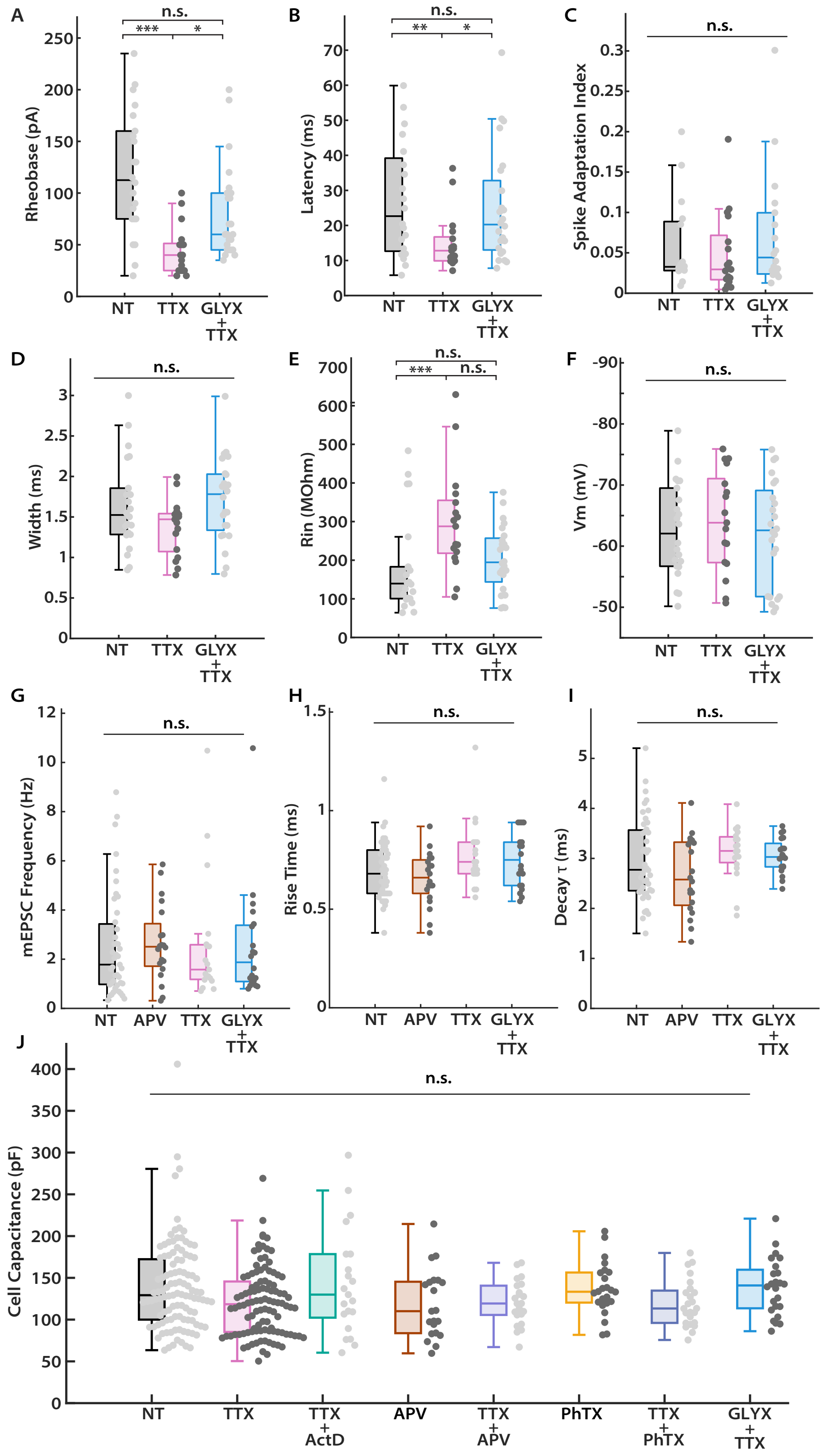
A
B
C
D
E
F
G
H
I
J
Rheobase (pA)
Latency (ms)
Spike Adaptation Index
Width (ms)
Rin (MOhm)
Vm (mV)
mEPSC Frequency (Hz)
Rise Time (ms)
Decay τ (ms)
Cell Capacitance (pF)
n.s.
***
**
*
NT
TTX
GLYX + TTX
APV
TTX + ActD
TTX + APV
PhTX
TTX + PhTX

Figure S5. NMDAR positive allosteric modulator GLYX does not affect synaptic and intrinsic cellular properties. Related to Figure 6.

(A) Comparison of rheobase current from the indicated conditions. Kruskal-Wallis test with Tukey correction: NT vs. TTX, p = 2.5E-5; NT vs. GLYX+TTX, p = 0.12; TTX vs. GLYX+TTX, p = 0.02.

(B) Comparison of latency to the first spike at 200 pA of current injection from the indicated conditions. Kruskal-Wallis test with Tukey correction: NT vs. TTX, p = 9.7E-3; NT vs. GLYX+TTX, p = 0.67; TTX vs. GLYX+TTX, p = 0.02.

(C) Comparison of spike frequency adaptation at 200 pA of current injection from the indicated conditions. Kruskal-Wallis test: p = 0.43.

(D) Comparison of full width at half-maximum for the first spike at rheobase from the indicated conditions. Kruskal-Wallis test: p = 0.13.

(E) Comparison of neuronal input resistance from the indicated conditions. Kruskal-Wallis test with Tukey correction: NT vs. TTX, p = 3.7E-3; NT vs. GLYX+TTX, p = 0.45; TTX vs. GLYX+TTX, p = 0.08.

(F) Comparison of resting membrane potential from the indicated conditions. Kruskal-Wallis test: p = 0.32.

(G) Comparison of the mean mEPSC frequency from the indicated conditions. Kruskal-Wallis test: p = 0.72.

(H) Comparison of the mEPSC rise time from the indicated conditions. Kruskal-Wallis test: p = 0.08.

(I) Comparison of the mEPSC decay time constant ($\tau$) from the indicated conditions. Kruskal-Wallis test: p = 0.14.

(J) Comparison of cell capacitance from the indicated conditions (all conditions examined for cultured neurons in the current study). Kruskal-Wallis test: p = 0.12.

A through F: same sample sizes as shown in Figure 6B.

G through I: same sample sizes as shown in Figure 6E.

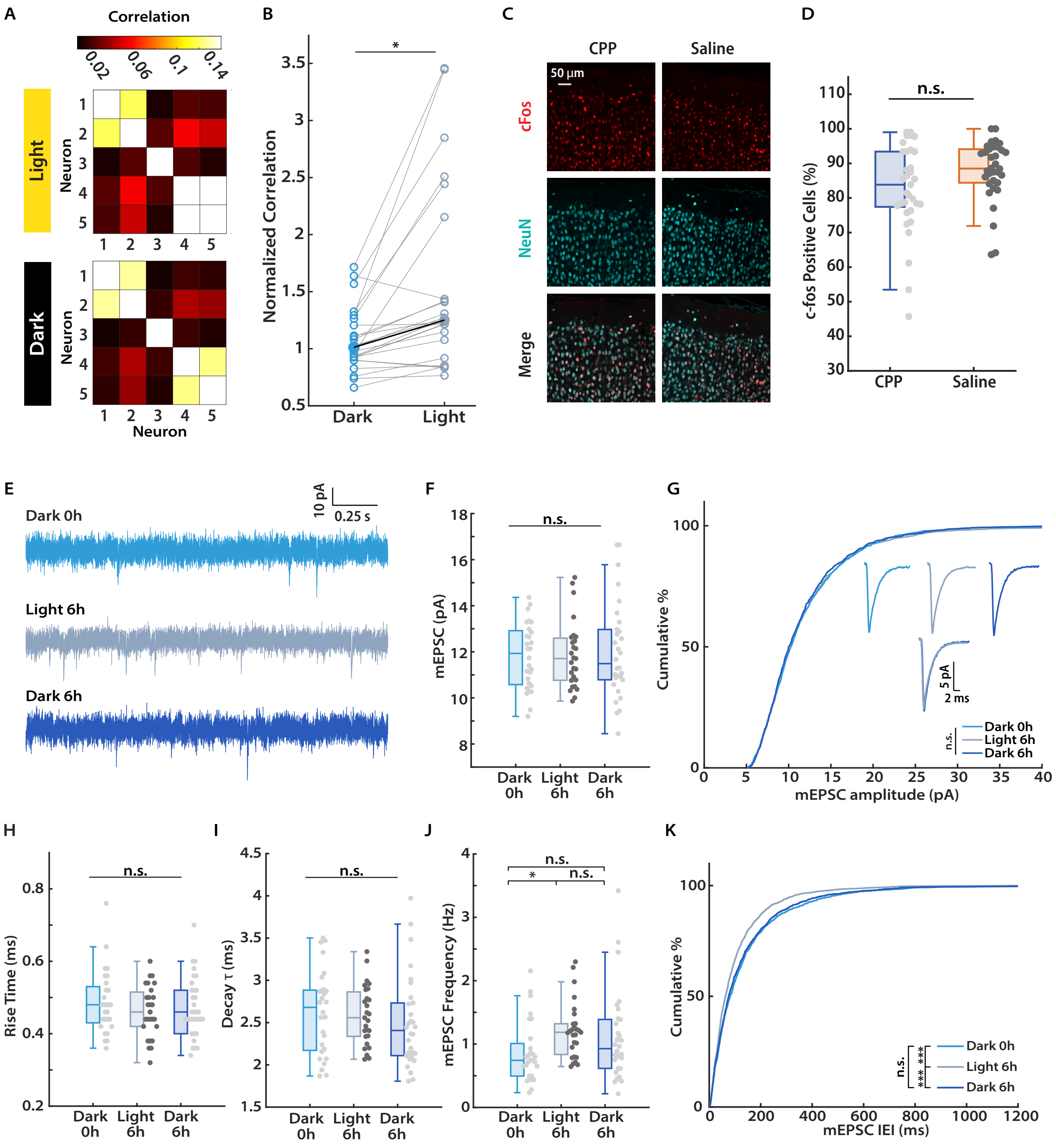
A
Correlation
0.02
0.06
0.1
0.14
Light
Dark
Neuron
B
Normalized Correlation
Dark
Light
C
CPP
Saline
50 µm
cFos
NeuN
Merge
D
n.s.
c-fos Positive Cells (%)
CPP
Saline
E
Dark 0h
Light 6h
Dark 6h
10 pA
0.25 s
F
n.s.
mEPSC (pA)
G
Cumulative %
mEPSC amplitude (pA)
5 pA
2 ms
Dark 0h
Light 6h
Dark 6h
H
Rise Time (ms)
I
Decay τ (ms)
J
mEPSC Frequency (Hz)
K
Cumulative %
mEPSC IEI (ms)

Figure S6. Light-driven increase in correlation does not affect synaptic strength in freely behaving animals. Related to Figure 7.

(A) Example pairwise correlation matrices of 5 neurons from 1 hemisphere during light (upper) and dark (lower), respectively. Each square in the matrix indicates the correlation value from a pair of neurons.

(B) Comparison of mean normalized correlation across light and dark for all pairs of neurons. Hollow circles connected with grey lines indicate individual pairs, whereas solid circles connected with a black line indicate the medians. Mann-Whitney U test, $p = 0.02$.

(C) Representative images of cFos expression in the L2/3 of mouse V1 following acute CPP (left) and saline (right) injections, respectively.

(D) Quantification and comparison of the percentage of cFos-positive neurons (out of all NeuN-positive neurons) following CPP and saline injections. Each data point indicates the cFos positive rate calculated from a region of interest in L2/3 selected from a sectioned V1-containing slice. CPP, n = 33, 3 animals; Saline, n = 34, 3 animals. Mann-Whitney U test: $p = 0.08$.

(E) Representative traces of mEPSC recordings in L2/3 pyramidal neurons from the three indicated timepoints in Figure 7E.

(F) Comparison of the mean mEPSC amplitude for L2/3 pyramidal neurons from the three indicated timepoints. Dark 0h, n = 28, 3 animals; Light 6h, n = 27, 3 animals; Dark 6h, n = 30, 3 animals. Kruskal-Wallis test: $p = 1.0$.

(G) Cumulative distribution of mEPSC amplitudes from the three indicated timepoints. Inset: Upper, unscaled average waveforms from the three timepoints, respectively; Lower, overlay of all three waveforms. Kolmogorov-Smirnov test: Dark 0h vs. Light 6h, $p = 0.90$; Dark 0h vs. Dark 6h, $p = 0.30$; Light 6h vs. Dark 6h, $p = 0.21$.

(H) Comparison of the mEPSC rise time from the three indicated timepoints. Kruskal-Wallis test: $p = 0.75$.

(I) Comparison of the mEPSC decay time constant ($\tau$) from the three indicated timepoints. Kruskal-Wallis test: $p = 0.34$.

(J) Comparison of the mean mEPSC frequency from the three indicated timepoints. Kruskal-Wallis test with Tukey correction: Dark 0h vs. Light 6h, $p = 0.01$; Dark 0h vs. Dark 6h, $p = 0.31$; Light 6h vs. Dark 6h, $p = 0.30$.

(K) Cumulative distribution of inter-event intervals (IEIs) from the three indicated timepoints. Kolmogorov-Smirnov test: Dark 0h vs. Light 6h, $p = 1.1E-12$; Dark 0h vs. Dark 6h, $p = 0.14$; Light 6h vs. Dark 6h, $p = 1.3E-8$.

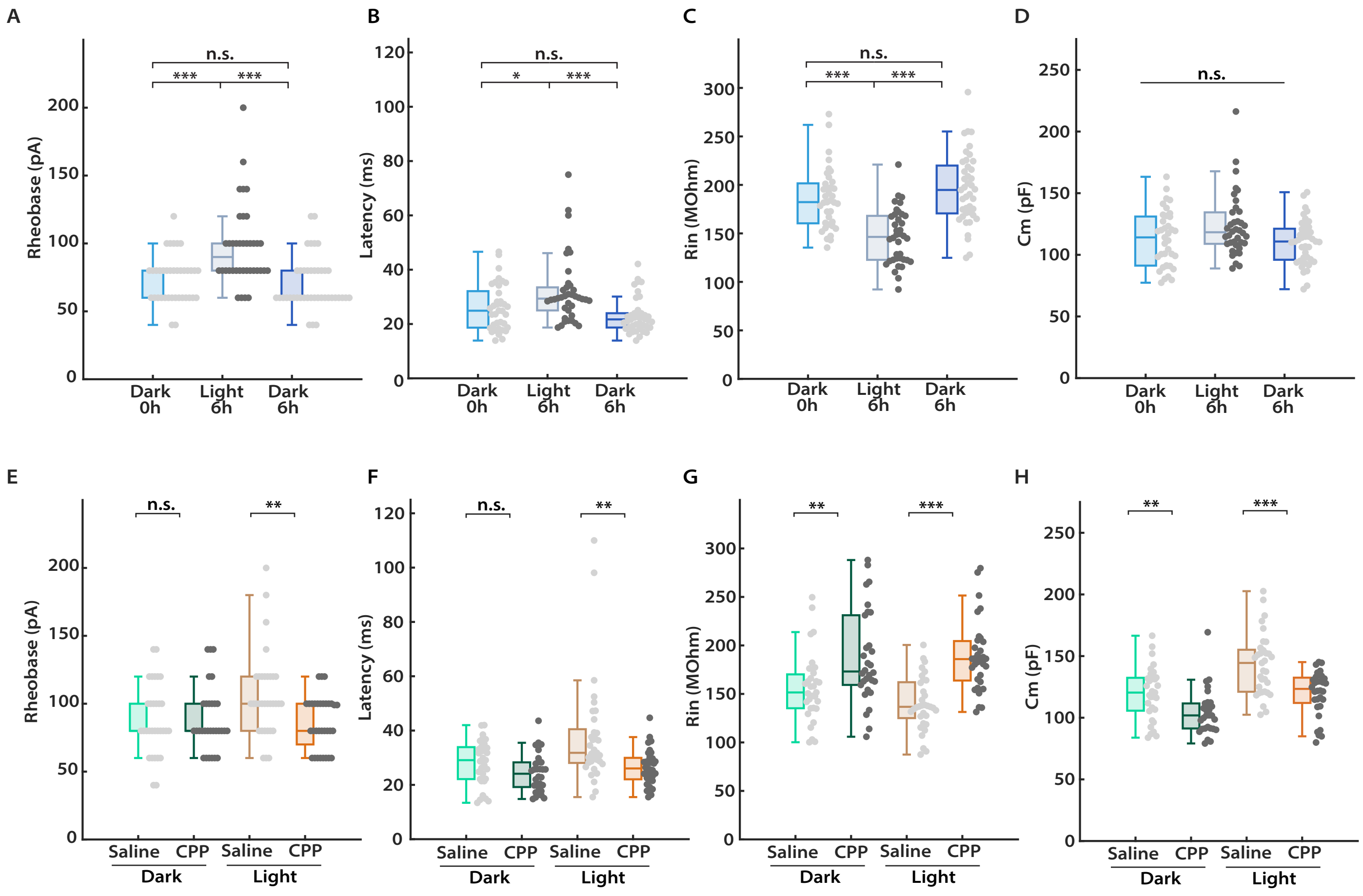
A
B
C
D
Rheobase (pA)
Latency (ms)
Rin (MOhm)
Cm (pF)
n.s.
***
*
Dark 0h
Light 6h
Dark 6h
E
F
G
H
**
Saline
CPP
Dark
Light

Figure S7. Light-driven IHP in freely behaving animals shares cellular changes with TTX- and DREADD-induced IHP. Related to Figure 7.

(A) Comparison of rheobase current from the indicated conditions. Kruskal-Wallis test with Tukey correction: Dark 0h vs. Light 6h, p = 6.0E-6; Dark 0h vs. Dark 6h, p = 0.54; Light 6h vs. Dark 6h, p = 3.3E-6.

(B) Comparison of latency to the first spike at 200 pA of current injection from the indicated conditions. Kruskal-Wallis test with Tukey correction: Dark 0h vs. Light 6h, p = 0.03; Dark 0h vs. Dark 6h, p = 0.28; Light 6h vs. Dark 6h, p = 1.2E-4.

(C) Comparison of neuronal input resistance from the indicated conditions. Kruskal-Wallis test with Tukey correction: Dark 0h vs. Light 6h, p = 1.2E-5; Dark 0h vs. Dark 6h, p = 0.52; Light 6h vs. Dark 6h, p = 1.5E-8.

(D) Comparison of cell capacitance from the indicated conditions. Kruskal-Wallis test: p = 0.08.

(E) Comparison of rheobase current from the indicated conditions. Mann-Whitney U test: Dark, CPP vs. Saline, p = 0.93; Light, CPP vs. Saline, p = 4.9E-3.

(F) Comparison of latency to the first spike at 200 pA of current injection from the indicated conditions. Mann-Whitney U test: Dark, CPP vs. Saline, p = 0.06; Light, CPP vs. Saline, p = 6.9E-4.

(G) Comparison of neuronal input resistance from the indicated conditions. Mann-Whitney U test: Dark, CPP vs. Saline, p = 4.8E-3; Light, CPP vs. Saline, p = 3.4E-7.

(H) Comparison of cell capacitance from the indicated conditions. Mann-Whitney U test: Dark, CPP vs. Saline, p = 3.5E-3; Light, CPP vs. Saline, p = 4.1E-4.

A through D: same sample sizes as shown in Figures 7C.

E through H: same sample sizes as shown in Figures 7F and 7I.

Table S1. Cell capacitance and resting membrane potential values for *ex vivo* slice experiments using DREADDs. Related to figures listed below.

| Experimental Conditions | Capacitance (pF) | Resting Membrane Potential (mV) |
|---|---|---|
| **Figure 2G**[a] | | |
| CNO + Saline | 59 ± 3 | -84 ± 1 |
| DR + CNO + Saline | 56 ± 3 | -85 ± 1 |
| **Figure 2I**[b] | | |
| CNO + Xpro | 57 ± 4 | -85 ± 1 |
| DR + CNO + Xpro | 57 ± 3 | -85 ± 1 |
| **Figure 4D**[c] | | |
| NT | 66 ± 4 | -80 ± 1 |
| CT+ | 64 ± 4 | -82 ± 1 |
| CT- | 61 ± 3 | -82 ± 1 |
| **Figure 6C**[d] | | |
| CNO + Saline | 64 ± 2 | -79 ± 1 |
| DR + CNO + Saline | 57 ± 2 | -80 ± 1 |
| CNO + CPP | 59 ± 2 | -78 ± 2 |
| DR + CNO + CPP | 59 ± 3 | -77 ± 1 |

a Capacitance ($C_m$): $p = 0.74$; Resting membrane potential ($V_m$): $p = 0.41$.

b $C_m$: $p = 0.96$; $V_m$: $p = 0.49$.

c $C_m$: $p = 0.52$; $V_m$: $p = 0.33$.

d $C_m$: $p = 0.17$; $V_m$: $p = 0.11$.
